# Distributed Representation of Taste Quality by Second-Order Gustatory Neurons in *Drosophila*

**DOI:** 10.1101/2020.11.10.377382

**Authors:** Nathaniel J. Snell, John D. Fisher, Griffin G. Hartmann, Mustafa Talay, Gilad Barnea

## Abstract

Sweet and bitter compounds excite different sensory cells and drive opposing behaviors. It is commonly thought that the neural circuits linking taste sensation to behavior conform to a labeled-line architecture, but in *Drosophila*, evidence for labeled lines beyond first-order neurons is lacking. To address this, we devised *trans-*Tango(activity), a strategy for calcium imaging of second-order gustatory projection neurons based on *trans*-Tango, a genetic transsynaptic tracing technique. We found distinct projection neuron populations that respond to sweet and bitter tastants. However, the bitter-responsive population was also activated by water alone. We further discovered that bitter tastants evoke activity upon both stimulus onset and offset. Bitter offset responses are exhibited by both first- and second-order gustatory neurons, but these responses are distributed among multiple types of projection neurons in the second order. These findings suggest a more complex coding scheme for gustatory information than can be explained by a labeled line model.

## INTRODUCTION

Taste enables animals to assess the nutritional value of food prior to ingestion. Sweet and bitter tastes respectively signal the presence of potentially calorie-rich sugars or toxic compounds. Accordingly, sweet tastants elicit food acceptance behaviors and bitter tastants elicit rejection, an innate behavioral response conserved between humans and the fly *Drosophila melanogaster* (Liman et al., 2014). Yet, despite the importance of taste in shaping feeding behavior, the neural circuits and codes linking taste sensation to behavioral response are not well understood.

The neural representation of taste begins with the detection of tastants by peripheral sensory cells. Humans can distinguish five taste qualities – sweet, bitter, salty, sour, and umami – and there are five distinct populations of taste bud cells, each tuned to one of these qualities (Yarmolinsky et al., 2009). In flies, the detection of sweet and bitter tastants is similarly performed by distinct populations of gustatory receptor neurons (GRNs) (Thorne et al., 2004; Wang et al., 2004; Marella et al. 2006). GRNs innervate hair-like sensilla on the tip of the proboscis, or labellum, where food intake occurs; they are also found in the pharynx and on the legs, wings, and ovipositor (Vosshall & Stocker, 2007). Sweet and bitter GRNs express different repertoires of gustatory receptors (GRs) (Clyne et al., 2000; Dunipace et al., 2001; Scott et al., 2001), endowing them with their tastant sensitivity (Dahanukar et al., 2001; Moon et al., 2006). Segregation of sweet and bitter GR expression enables genetic access to these populations: Gr64f and Gr66a are expressed in all sweet and bitter GRNs of the labellum, respectively (Dahanukar et al., 2007; Weiss et al., 2011). GRN axons project to a region of the brain called the subesophageal zone (SEZ), and intriguingly, axons of sweet and bitter GRNs terminate in different areas of the SEZ, suggesting that they may synapse onto different downstream circuits (Thorne et al. 2004; Wang et al., 2004).

Two predominant models have been proposed for the circuit logic that underlies taste information processing from the peripheral taste cells to higher brain regions (Yarmolinsky et al., 2009; Simon et al., 2006). The labeled line model posits that taste information is relayed through parallel, segregated channels from the periphery to central brain. By contrast, the distributed coding model holds that different tastes activate overlapping neuronal ensembles, which are read out by downstream brain areas as a population code. The existence of separate sweet and bitter GRN populations with distinct projection patterns within the SEZ is consistent with a labeled line model (Thorne et al., 2004; Wang et al., 2004). Labeled line coding in peripheral neurons is further supported by behavioral evidence. Optogenetically activating sweet GRNs is sufficient to drive proboscis extension and feeding behaviors, while silencing or ablating these neurons strongly suppresses these behaviors; likewise, activation of bitter GRNs prevents feeding, while silencing or ablating these neurons promotes proboscis extension and feeding responses to sweet tastants even in the presence of aversive bitter compounds (Thorne et al., 2004; Wang et al., 2004; Moreira et al., 2019; Musso et al., 2019). Moreover, several putative second-order gustatory neurons respond to either sweet or bitter, but not both (Kain and Dahanukar 2015; Yapici et al. 2015; Miyazaki et al. 2016; Kim et al. 2017; Bohra et al. 2018). In other insect species, however, recordings from second-order neurons have revealed complex tuning properties that seem inconsistent with labeled lines (Kvello et al. 2010; Reiter et al. 2015). Beyond the second order of the circuit, data suggests that sweet and bitter activate separate central neuronal populations (Harris et al. 2015). Yet, since these tastes evoke different behavioral responses, at some point in the circuit the responding neuronal populations must be different. Determining which model is operative therefore requires understanding the tuning of gustatory neurons at each successive stage in the circuit.

We recently developed *trans*-Tango, a genetic technique for transsynaptic tracing and circuit manipulation, and used it to identify several types of second-order gustatory neurons that are postsynaptic to sweet GRNs (Talay et al. 2017). The transsynaptic output of *trans*-Tango is transcription of a marker protein; by changing the identity of this protein, *trans*-Tango can be used for purposes beyond anatomical tracing. For example, using a genetically encoded calcium indicator as the transsynaptic reporter would enable one to measure neural activity across a population of second-order neurons in a circuit.

Here, we establish a configuration of *trans*-Tango enabling *in vivo* calcium imaging of transsynaptically labeled neurons. We use this strategy to examine the taste coding models by determining how sweet and bitter are encoded by second-order gustatory projection neurons (GPNs) in *Drosophila*. We find that sweet and bitter tastants excite different GPNs innervating distinct regions of the superior protocerebrum. These GPNs respond similarly to stimuli of the same taste quality, although for sweet tastants, the degree of this similarity is modulated by hunger state. While these findings are consistent with a labeled line model, we report that water also drives activity across the bitter-responsive region of GPNs. Although bitter taste and water thus activate similar GPNs, we discover a property of taste coding that may distinguish these stimuli: bitter tastants alone activate GPNs upon both their presentation (stimulus onset) and removal (stimulus offset). This bitter offset response originates in the first-order GRNs encoding bitter onset, but in second-order GPNs, bitter offset activity is distributed beyond the bitter onset-responsive region. Our results support a mixed model of taste coding, wherein some neurons are selective for a single taste quality while distributed populations are recruited to represent others.

## RESULTS

### Anatomy of Second-Order Neurons in the Sweet and Bitter Circuits

Sweet and bitter tastants activate separate classes of GRNs, but it is not clear whether this labeled line representation is maintained beyond the first order of the circuit. If labeled lines are maintained, one possibility is an extreme case, wherein the second-order neurons of the sweet and bitter circuits are completely distinct. To test this, we used *trans*-Tango to compare the neural populations postsynaptic to *Gr64f*+ sweet GRNs and *Gr66a*+ bitter GRNs. Surprisingly, similar patterns of labeling were observed in both conditions, including dense innervation of the SEZ and multiple tracts projecting to the superior neuropils, to which we refer here as the lateral, mediolateral, and medial SEZ tracts (lSEZt, mlSEZt, mSEZt) (Talay et al., 2017; Figures 1A-1B). While the overall labeling from each genotype is similar, there may be subtle distinctions. For example, *Gr66a*>*trans*-Tango appears to label the mlSEZt more strongly than the lSEZt, while *Gr64f*>*trans*-Tango labels both tracts at comparable levels. Nevertheless, the morphology of projection terminals in and around the superior lateral protocerebrum (SLP) is similar in these two strains (Figure 1D). It is unlikely that completely distinct populations are labeled in these two strains, since driving *trans*-Tango from both Gal4 drivers simultaneously did not appear to label a substantially greater number of neurons (Figure 1C). However, from these results we cannot conclude the degree to which these second-order neuron populations overlap.

**Figure 1.**
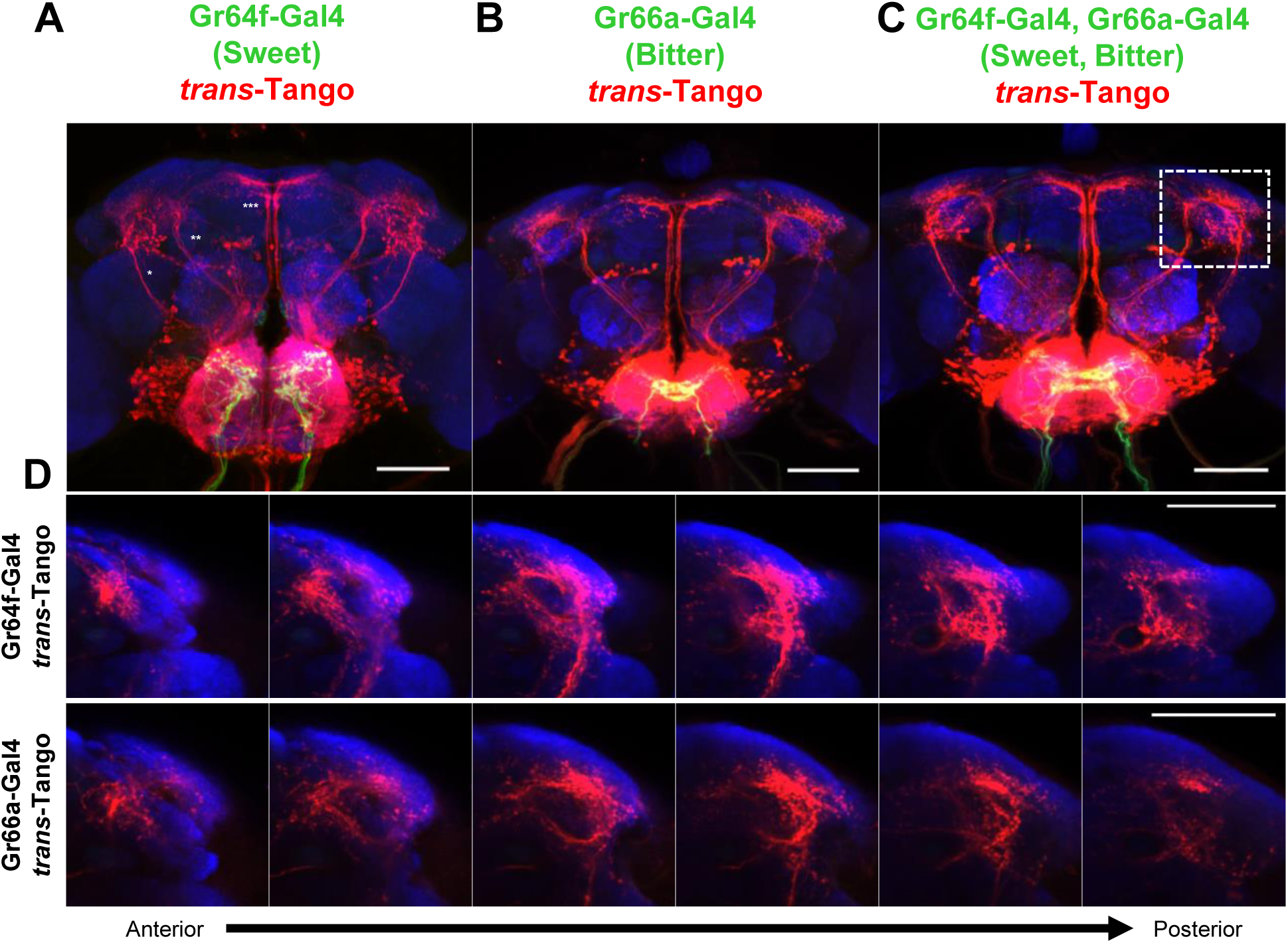
Anatomy of the second-order gustatory neurons. (A) Anatomy of second-order gustatory neurons postsynaptic to sweet GRNs, as revealed by *Gr64f-Gal4* > *trans*-Tango, with three SEZ tracts labeled: * = lSEZt; ** = mlSEZt; *** = mSEZt. (B) Anatomy of second-order gustatory neurons postsynaptic to bitter GRNs, as revealed by *Gr66a-Gal4* > *trans*-Tango. (C) Anatomy of second-order gustatory neurons postsynaptic to both sweet and bitter GRNs, revealed by *Gr64f-Gal4, Gr66a-Gal4* > *trans*-Tango. (D) Projection neurons targeting SLP labeled by *trans*-Tango initiated by either *Gr64f-Gal4* or Gr66a-Gal4 have similar overall morphology. Images correspond to area within white dotted line in C; sections from left to right depict progressively more posterior planes of depth. In all panels, first-order neurons are labeled by GFP (green) and second-order neurons by tdTomato (red). Scalebars in all panels 50 μm; for D, scalebar in rightmost image applies to entire row.

### Sweet and Bitter Stimuli Activate Different GPNs

Understanding the neural representation of taste requires determining not just the anatomy of relevant neurons, but also their tuning properties. Thus, we decided to perform *in vivo* calcium imaging of second-order gustatory neurons using *trans*-Tango, a strategy we term *trans*-Tango(activity). The modular design of *trans*-Tango enables the expression of any reporter protein, not just a fluorescent marker, in the second-order neurons of a circuit. The combination of cell-type-specific Gal4, *trans*-Tango, and a calcium indicator under QUAS control as the readout of *trans*-Tango would allow calcium imaging of the neurons postsynaptic to the Gal4+ starter neurons. However, calcium imaging of neuropil requires high levels of calcium indicator expression, and *trans*-Tango reporter levels can take weeks to accumulate to saturation (Talay et al., 2017). To overcome this challenge, we generated new QUAS-GCaMP6 reagents with significantly boosted expression levels (see details in Methods). As an output of *trans*-Tango, these reporters enabled robust *in vivo* calcium imaging of second-order neurons in flies as young as 10 days post-eclosion.

Previously, we used mosaic analysis to identify several classes of second-order GPNs that comprise the three SEZ tracts as well as local interneurons of the SEZ (Talay et al., 2017). Our objective here is to understand how taste is represented in the circuits connecting peripheral neurons to central brain regions, so we limited our investigation to GPNs rather than local interneurons. To simultaneously image all GPNs of the sweet and bitter circuits, we expressed GCaMP6s under control of *trans-*Tango initiated from both *Gr64f-Gal4* and *Gr66a-Gal4* together. We recorded calcium responses of GPNs to the presentation of tastants to the labellum. We imaged the neuropil of GPN arborizations in the SLP, a region previously implicated in innate behaviors. Because GPNs targeting the SLP project to several planes of depth along the anterior-posterior axis, we imaged a volume of several planes that captured the full population.

Two sweet tastants (maltose and sucrose) and two bitter tastants (denatonium and papaverine) all evoked strong activity in GPNs, but the calcium responses to these tastants were distributed unevenly among the GPN terminals. Both maltose and sucrose evoked activity most prominently in dorsal and lateral regions within the anterior planes of the volume, while both denatonium and papaverine drove activity in more ventral and medial posterior areas (Figure 2A and 2C). Thus, patterns of GPN activity were similar within a taste quality, but dissimilar between qualities. These response patterns were consistent across multiple trials and across flies (Figure S1A-B). Correlations between responses further supported the separation of sweet and bitter representations: GPN responses to tastants of the same quality were highly correlated, but responses to tastants of different qualities were not (Figure 2B and S1D).

**Figure 2.**
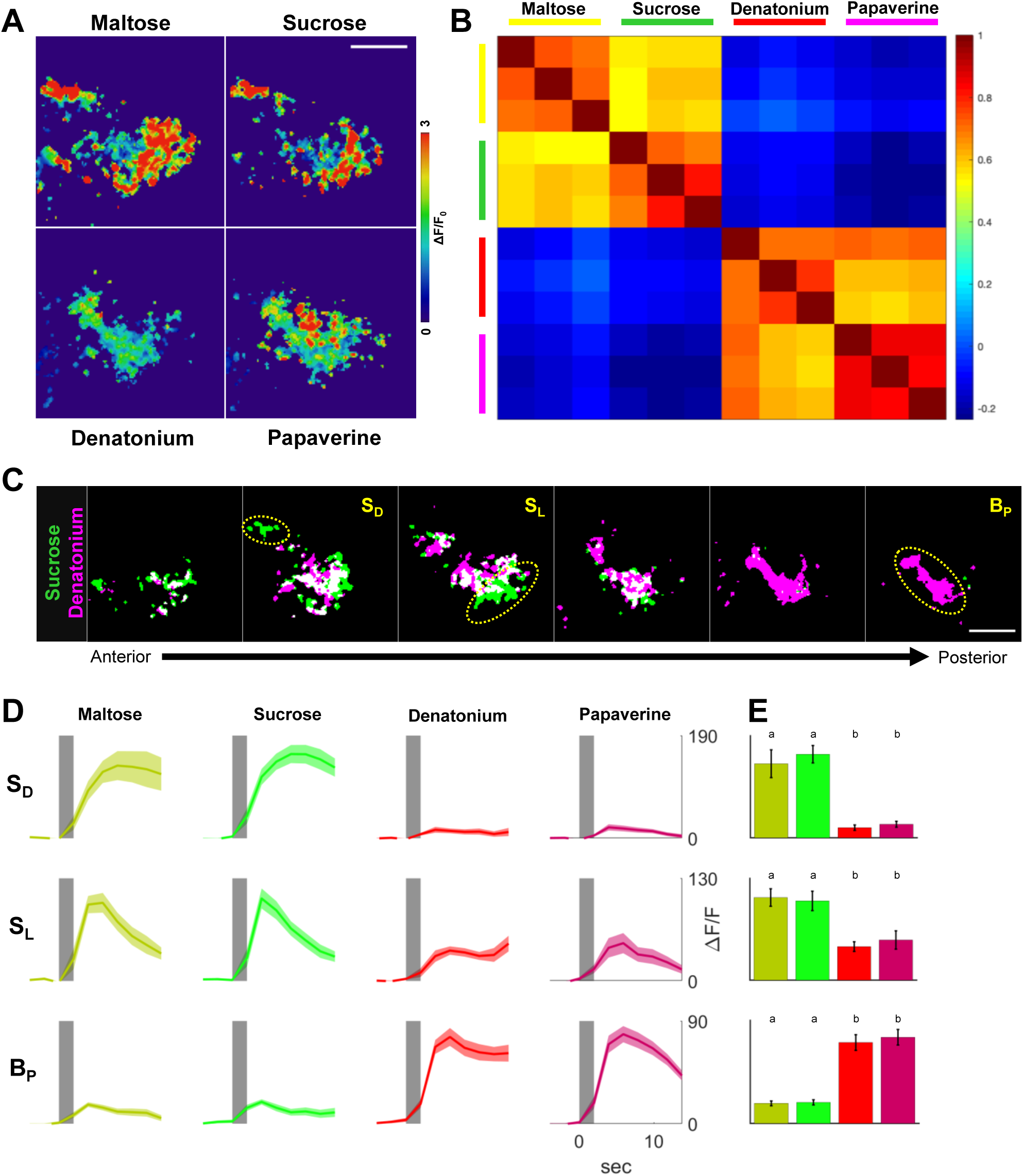
Sweet and bitter tastants excite different populations of GPNs. (A) Representative heatmaps of GPN responses to sweet (maltose, sucrose) and bitter (denatonium, papaverine) tastants. Heatmaps are maximum projections over all planes in the volume; each image represents the response during one trial, and all images are from the same experiment. Heatmaps show ΔF/F value from 0 to 3 as indicated by legend on the right. (B) Pairwise peak response correlations (Pearson’s r) between trials, group-averaged (n = 11 flies). Each group of three rows/columns represents three consecutive trials of the tastant indicated above matrix. Colors represent correlation coefficient value according to legend on the right. (C) Representative images of spatial patterns of significant GPN responses to sucrose (green) and denatonium (magenta) in individual planes. Images are ordered from anterior to posterior; white indicates overlap between areas of significant response to each tastant. Yellow dotted ovals indicate locations of ROIs S_L_, S_D_, and B_P_. (D) Fluorescence traces showing magnitude and time course of responses within each ROI. Responses are color-coded by tastant as follows: maltose, yellow; sucrose, green; denatonium, red; papaverine, magenta. Bold trace denotes mean and shaded region denotes SEM. Gray bar denotes time of stimulus onset. Y axis is ΔF/F; scale is same for panels within a row. X axis is time; scale is same for all panels. (E) Peak responses to each tastant within each ROI. Colors represent tastants with same color scheme as D. Y axis is maximum ΔF/F; scale is same as 2D. Error bars denotes SEM. Letters represent statistically significant differences between groups (p < .05, one-way ANOVA with Tukey post-hoc test). All scalebars 30 μm; scalebar applies to all images in a panel. See also Figures S1-S2.

While sweet and bitter responses were distinct at the population level, we considered the possibility that a subset of the GPN population might respond to both qualities. To identify whether any such regions were present in our volume, we determined the spatial patterns of the activity evoked by each tastant within each plane of depth, and assessed the degree of overlap between the patterns for different tastants. Minimal overlap was observed between sweet and bitter regions in any single plane (Figure 2C). In contrast, tastants of the same quality often activated nearly completely overlapping regions (Figure S1E). This trend held across flies: most pixels, but not all, responded selectively to only one taste quality in every fly tested (Figure S1C).

Typically, the pixels that showed significant responses to both taste qualities did not cluster together in space, but rather were scattered within areas selective for just one quality. We wondered whether these pixels represented weak responses to the opposite taste quality or just noise. To investigate this, we used the spatial response patterns we derived for each tastant to delineate ROIs in single planes that corresponded to different sweet-responsive or bitter-responsive regions of the volume (Figure 2C). We considered two ROIs in the anterior sweet-responsive region: one corresponding to a projection along the dorsal edge of the region (S_D_), and another corresponding to a region along the lateral edge (S_L_). While these two regions are located near each other, they likely comprise separate GPNs, as the time courses of their responses were distinct: the time to peak response was shorter for S_L_ than S_D_, and the sweet response in S_L_ adapted faster than the response in S_D_ (Figure 2D). Furthermore, while S_D_ was selective for sweet tastants, S_L_ responded to both sweet and bitter tastants, though the magnitude of the bitter responses was lower (Figure 2D). We performed a similar analysis for the bitter-responsive region, defining ROIs within this region at three levels of depth within the volume. We refer to these as the anterior (B_A_), intermediate (B_I_), and posterior (B_P_) bitter areas, and based the demarcation of these ROIs on stereotyped morphology of the bitter-responsive neuropil to ensure consistency across individual flies (Figure S1E). Within each of these ROIs, both bitter tastants elicited strong activation at comparable levels. While both sweet tastants also produced minor responses within each of these regions, these were significantly weaker than the bitter-evoked responses (Figure 2E). Thus, each GPN region responded primarily to only one of the two taste qualities.

We considered the possibility that sweet GPNs might be specifically labeled by *trans*-Tango initiated from the sweet GRN Gal4 driver, and likewise for bitter GPNs and the bitter GRN driver. To test this possibility, we expressed GCaMP6s in the GPNs labeled in each of these strains and imaged their responses to sweet and bitter tastants. Surprisingly, whether we used either the sweet driver or the bitter driver, we observed responses to both sweet and bitter tastants in the GPNs labeled by *trans*-Tango (Figure S2A-D). However, the GPN region responsive to sweet tastants was markedly diminished in the bitter Gal4-driven *trans*-Tango flies, as was the region responsive to bitter tastants in the sweet Gal4-driven *trans*-Tango flies (Figure S2E-F). This result is consistent with different but overlapping subsets of GPNs that are labeled in the two cases, with each GRN type preferentially targeting GPNs tuned to the same taste quality.

### Water and Bitter Stimuli Activate Similar GPNs

Apart from sweet and bitter GRNs, the labellum contains both water-sensing GRNs and mechanoreceptors, so we next asked if mechanical stimulation or osmolarity contributed to the responses that we observed. To assess whether this was the case, we compared GPN responses to sucrose and denatonium to those elicited by water and polyethylene glycol (PEG; 3,350 g/mol; 20% w/v), a stimulus that does not excite water-sensing GRNs (Cameron et al., 2010). Both stimuli evoked GPN activity, indicating that these neurons are not activated strictly by sweet and bitter taste (Figure 3A). Moreover, the water and PEG response was highly correlated, suggesting that this water-evoked activity may largely be a mechanosensory response (Figure 3B and S3B).

**Figure 3.**
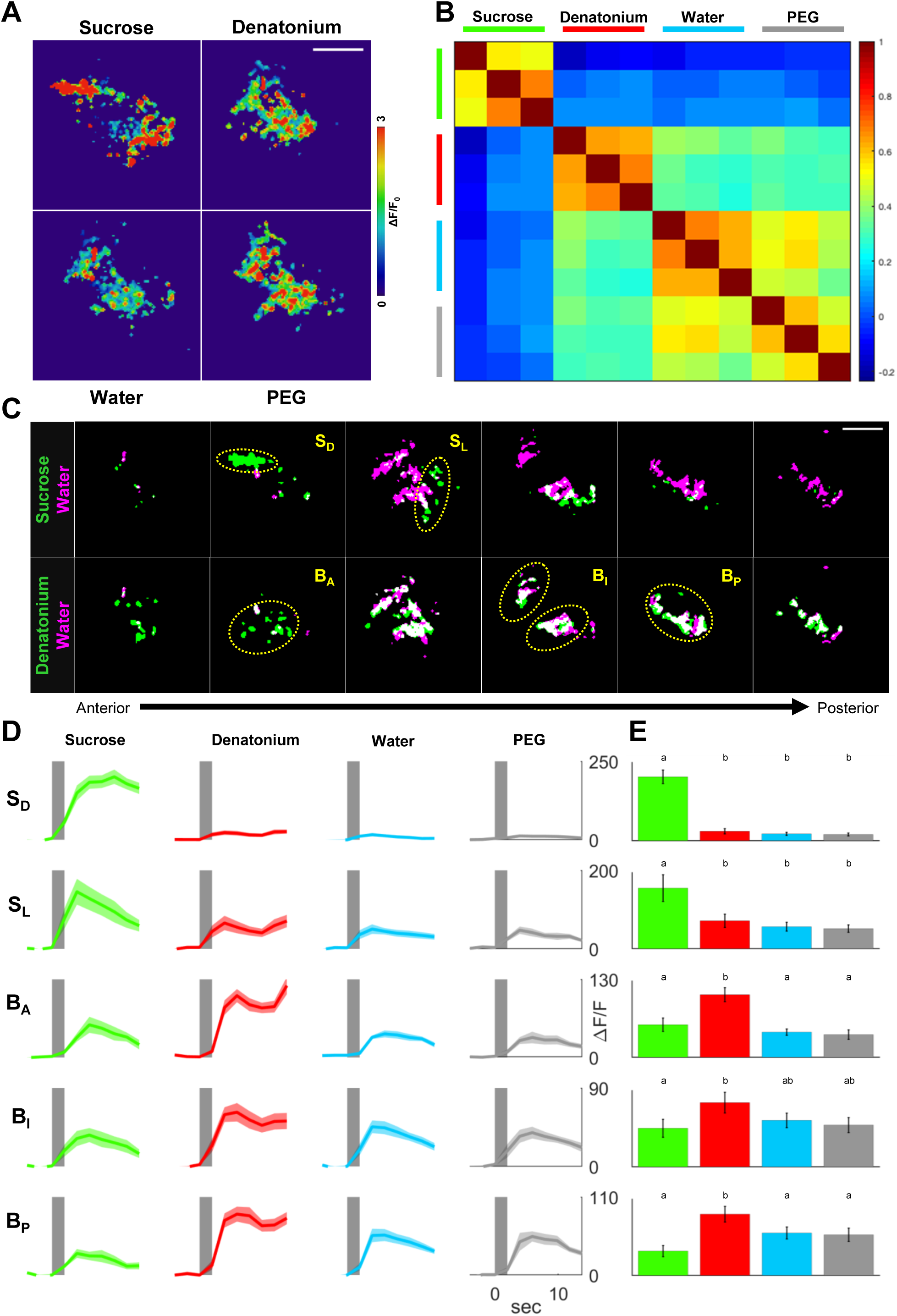
GPNs responding to bitter also respond to water and PEG. (A) Representative heatmaps of GPN responses to sucrose, denatonium, water, and PEG. Heatmaps are maximum projections over all planes in the volume; each image represents the response during one trial, and all images are from the same experiment. Heatmaps show ΔF/F value from 0 to 3 as indicated by legend on the right. (B) Pairwise peak response correlations (Pearson’s r) between trials, group-averaged (n = 8 flies). (C) Representative images of spatial response patterns to water (magenta) and either sucrose (green, top row) or denatonium (green, bottom row) in individual planes. Yellow dotted ovals indicate locations of ROIs S_D_, S_L_, B_A_, B_I_, and B_P_. (D) Fluorescence traces showing magnitude and time course of responses within each ROI. Responses are color-coded by tastant as follows: sucrose, green; denatonium, red; water, cyan; PEG, light gray. (E) Peak responses to each tastant within each ROI. Colors represent tastants with same color scheme as D. Y axis is maximum ΔF/F; scale is same as D. Error bars denote SEM. Letters represent statistically significant differences between groups (p < .05, one-way ANOVA with Tukey post-hoc test). All scalebars 30 μm; scalebar applies to all images in a panel. See also Figure S3.

Intriguingly, GPN responses to water and PEG also exhibited high correlation with the denatonium response, but no correlation with the sucrose response (Figure 3B and S3B). The spatial patterns of these responses reflected these correlations: water-responsive regions had minimal overlap with sucrose-responsive regions, but substantial overlap with areas responding to PEG and denatonium (Figure 3C and S3A). In each of the three ROIs in the bitter-responsive region, activity was observed in response to both water and PEG. However, denatonium drove stronger responses in these areas than either water or PEG; this difference in magnitude was largest in area B_A_, but was observed in areas B_I_ and B_P_ as well. Interestingly, the response to sucrose in each of these ROIs was comparable in magnitude to the water and PEG responses, indicating that the sucrose-driven activity in these regions may not be caused by the sweetness of the stimulus (Figure 3D-E). In contrast, neither water nor PEG produced significant activation of area S_D_, suggesting that this region selectively responds to sweet tastants. Water and PEG did elicit activity within area S_L_, but the magnitude of this activity was much smaller than sucrose-evoked activity, and comparable to denatonium-evoked activity.

### GPNs Respond Similarly to Multiple Tastants of the Same Taste Quality

While our initial experiments demonstrate that GPNs respond similarly to two tastants of the same quality, this pattern may not generalize. Indeed, in some insects, different sweet tastants evoke substantially different patterns of neural activity (Reiter et al., 2015). To test this, we imaged GPN calcium responses to a panel of five sweet tastants: fructose, glucose, maltose, sucrose, and trehalose, and included water as a control. Each of these stimuli elicited a response in GPNs (Figure 4A). However, there was substantial heterogeneity in the spatial patterns of these responses. Tastants produced responses that generally matched one of two spatial patterns: one in which activity was strongest in areas S_D_ and S_L_, and another in which activity was predominantly observed in more medial, water-responsive regions. As such, population-level responses to some sweet tastants were correlated with the water response (Figure 4C). These tastants did not exclusively activate the water-responsive region, as excluding the water-responsive pixels revealed substantial activity outside this area for all sweet tastants (Figure S4A). The heterogeneity in the response patterns of GPNs to the different sweet tastants was even more apparent after removing water-responsive pixels from the analysis (Figure S4C). Furthermore, the magnitude of responses in areas S_D_ and S_L_ varied substantially between stimuli (Figure S5A-B).

**Figure 4.**
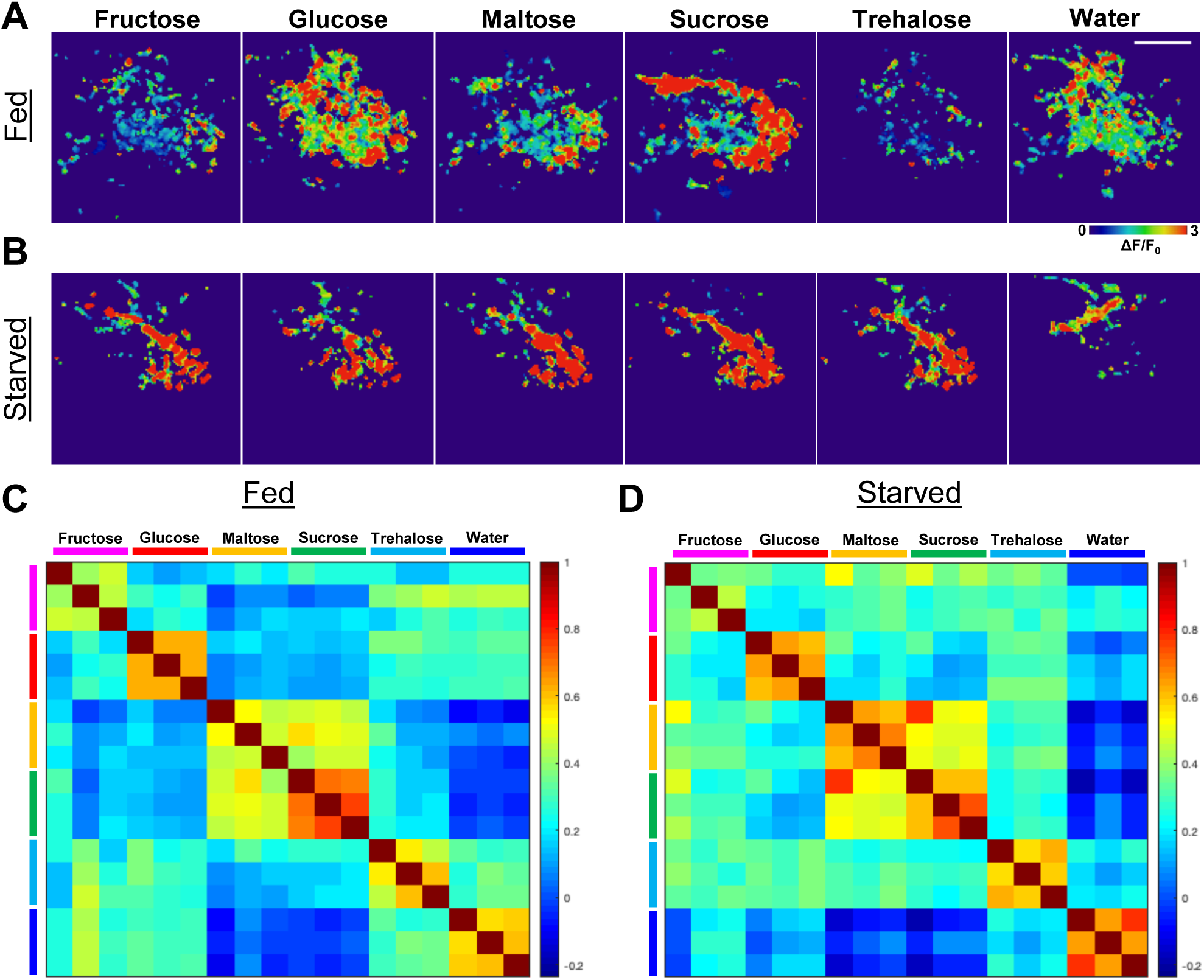
GPN responses to sweet vary between tastants, but are more consistent after starvation. (A) Heatmaps from a fed fly showing responses to a panel of sweet tastants. Heatmaps are maximum projections over all planes in the volume; each image represents the response during one trial, and all images are from the same experiment. Heatmaps show ΔF/F value from 0 to 3 as indicated by legend on the right. (B) Heatmaps as in A, but for a fly wet-starved for 1 day. (C) Matrix of inter-trial correlations for fed flies, group averaged (n = 8 flies). (D) Matrix of inter-trial correlations for starved flies, group averaged (n = 8 flies). Scalebar marks 30 μm and applies throughout figure. See also Figure S4-S5.

While the heterogeneity of GPN responses to sweet tastants might indicate distinct spatial representations of different sugars, there are other possible explanations. In particular, maltose and sucrose may drive more robust activity in sweet-responsive areas because these sugars are more potent activators of GRNs than most others (Dahanukar et al., 2007), and perhaps a threshold of GRN activity must be reached to robustly drive activity in sweet-responsive GPNs. Since starvation potentiates both sweet GRNs and some sweet-responsive second-order gustatory neurons (Inagaki et al., 2012; Kain & Dahanukar, 2015), we wondered if starved flies would exhibit stronger GPN responses to sweet tastants. Indeed, flies starved for 24 hours exhibited equal or stronger correlations between GPN responses for every pair of sweet tastants (Figure 4B and 4D and S5C). Excluding water-responsive pixels from the analysis enhanced this effect, both in the visibility of these responses (Figure S4B) and in the inter-trial correlations (Figure S4D). Correspondingly, responses to sweet tastants in areas S_D_ and S_L_ were equal or stronger in starved flies than in fed flies (Figure S5A-B).

We performed the equivalent experiments with a panel of bitter tastants: caffeine, denatonium, lobeline, papaverine, and quinine, with water included again as a control. As was the case for sweet compounds, each of these stimuli evoked activity in GPNs (Figure 5A). However, unlike for sweet tastants, GPN responses to bitter tastants were highly correlated with one another (Figure 5B and S6C). Moreover, all bitter responses (except papaverine) were significantly correlated with the water response, as was previously observed for denatonium. The spatial pattern of responses to each bitter stimulus nearly completely overlapped the pattern of the water response across each plane of the volume (Figure 5C). However, bitter tastants generally drove stronger responses than water in this region. This pattern was most pronounced in area B_A_, but was observed across planes of depth (Figure S6A-B). These results indicate that the spatial overlap between denatonium and water responses generalizes to bitter tastants as a group.

**Figure 5.**
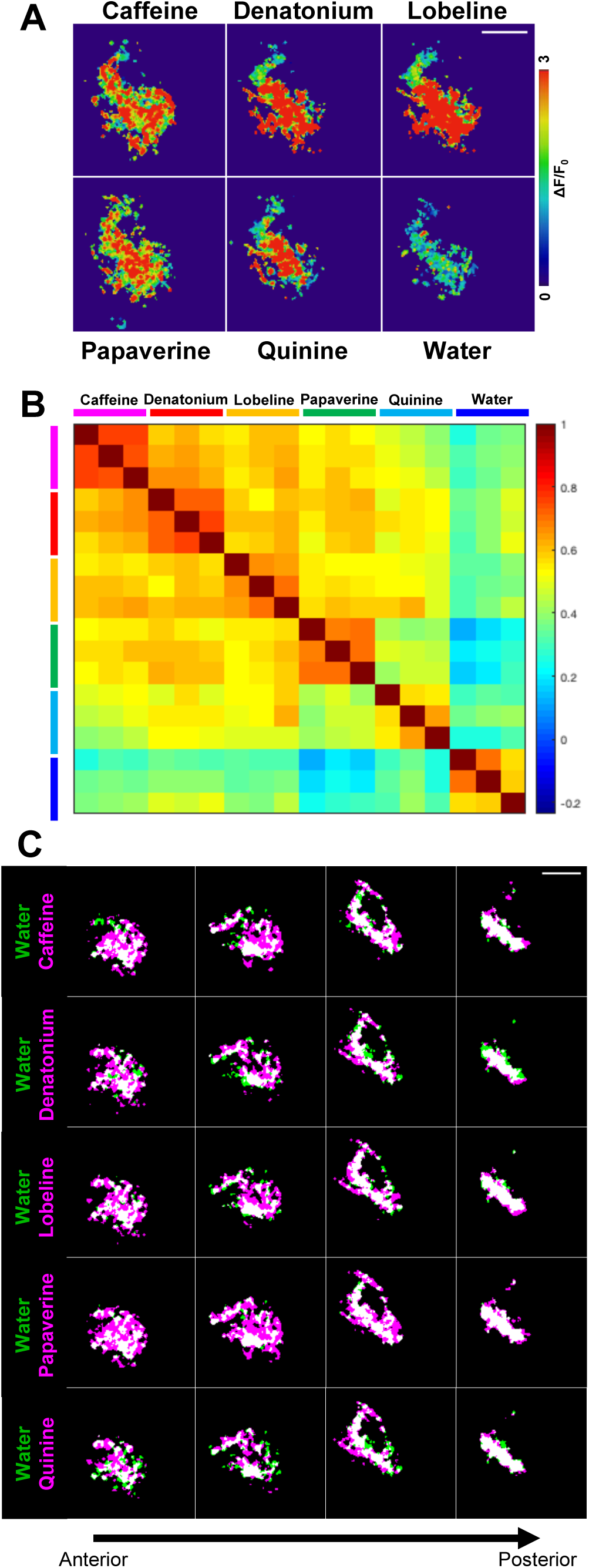
GPN responses to bitter are similar between tastants as well as to water. (A) Heatmaps showing peak responses to five bitter tastants and water. Heatmaps are maximum projections over all planes in the volume; each image represents the response during one trial, and all images are from the same experiment. Heatmaps show ΔF/F value from 0 to 3 as indicated by legend on the right. (B) Matrix of inter-trial correlations, group averaged (n = 8). (C) Spatial patterns of bitter-responsive areas (magenta) overlaid with the water-responsive area (green); white denotes overlap. All scalebars 30 μm; scalebar applies to all images in a panel. See also Figure S6.

### GRNs and GPNs Respond to Bitter Stimulus Offset in a Concentration-Dependent Manner

The finding that bitter tastants and water evoke responses within similar GPN regions raises the question of how these signals might be distinguished by downstream neurons. Because the observed magnitude of the bitter response was typically greater than that of water (Figure S6B), it is possible that relative activity levels are used to differentiate the two types of stimulus. However, this difference in magnitude was rather small for some bitter tastants, even though the concentrations we used are known to be highly aversive (Sellier et al., 2011; Weiss et al., 2011). Thus, the magnitude of GPN response alone is unlikely to provide enough information to reliably discriminate bitter from water, a critical ability for survival.

Interestingly, we observed that denatonium evoked a GPN response upon both the delivery of the stimulus (stimulus onset) and its removal (stimulus offset) (Figure 6A, left). This offset response was time-locked to the removal of the stimulus: when we held the bitter stimulus on the labellum for varying lengths of time, the second activity peak consistently followed stimulus offset, rather than a fixed time delay from the onset (Figure 6A, right). While responses to stimulus offset have been observed in the moth *Manduca sexta* (Reiter et al. 2015), no such effect has been reported in *Drosophila*. We wondered whether this effect emerged at the second order of the circuit or if it was initiated in the GRNs. To test this, we performed similar experiments while imaging calcium activity in bitter GRNs. We observed an offset response to denatonium in these first-order neurons as well (Figure 6B). Besides denatonium, we observed offset responses to lobeline and quinine in both GRNs and GPNs, suggesting that this offset response is a general property of bitter representation (Figure S7A-B). Furthermore, this response is specific to bitter; no offset response to water or sucrose was observed at either stage of the circuit (Figures 6C, 7A and data not shown).

**Figure 6.**
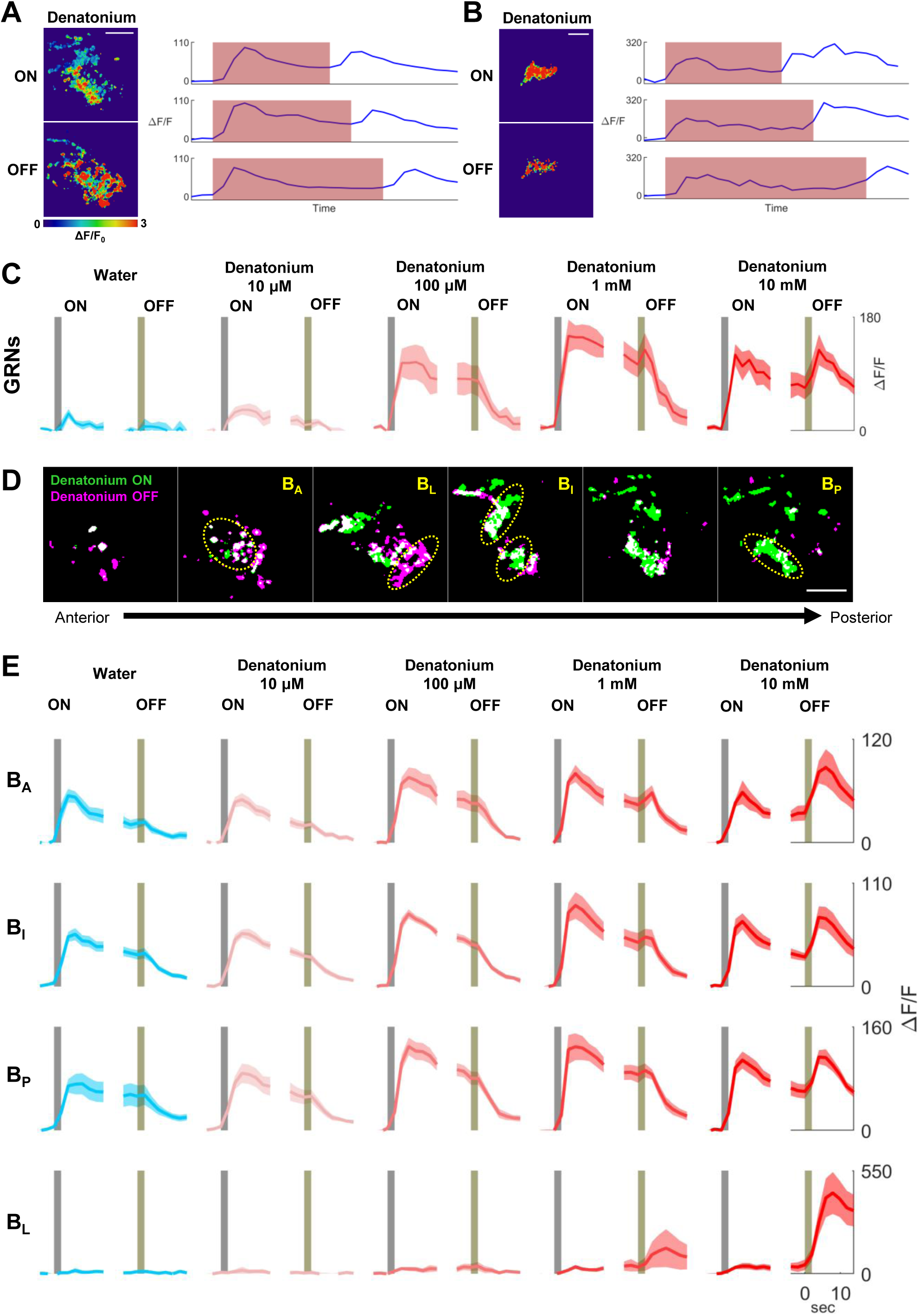
GRNs and GPNs respond to bitter stimulus offset. (A) GPN response to denatonium onset and offset. Left, representative heatmaps of onset and offset responses. Right, traces showing response to denatonium applied for different lengths of time. In traces, red shading denotes the time interval during which the stimulus was applied. Top row, time is approximately 20 sec; middle row, 25 sec; bottom row, 30 sec. (B) Same as A, but for Gr66a+ GRNs. (C) Gr66a+ GRN responses to increasing concentrations of denatonium. For each concentration, left plot shows responses aligned to stimulus onset (gray bar) and right plot shows responses aligned to offset (yellow bar). Plots show mean (bold trace) and SEM (shaded area; n = 5 flies, 3 trials per concentration). (D) Spatial maps of onset (green) and offset (magenta) responses to denatonium in GPNs across individual planes. Yellow dotted circles denote ROIs B_A_, B_L_, B_I_, and B_P_. (E) GPN responses to increasing concentrations of denatonium in four ROIs depicted in Figure D. Color scheme follows that of panel C (n = 6 flies, 3 trials per concentration). All scalebars 30 μm; scalebar applies to all images in a panel. See also Figure S7.

The concentrations of bitter tastants we initially used to evoke the offset responses were relatively high (10 mM for denatonium and lobeline, 100 mM for quinine). We next tested whether the concentration threshold for the offset response matched that of the onset response. GRNs exhibited a significant response to bitter onset at concentrations as low as 100 μM for denatonium (p<.05, two-sample one-tailed t-test with Bonferroni correction, denatonium response greater than water response, see Methods; Figure 6C). However, at this concentration, no offset response was clearly visible. A slight albeit visible activity peak at stimulus offset was detected at 1 mM, and a strong and significant offset response was seen at 10 mM (p<.05, two-sample one-tailed t-test with Bonferroni correction, denatonium response greater than water response; Figure 6E). The concentration thresholds obtained for onset and offset responses to lobeline were the same as for denatonium (Figure S7C).

We next sought to examine whether the responses to bitter onset and offset were similarly concentration-dependent in GPNs. We performed the equivalent set of experiments, imaging GPN activity to determine thresholds for the bitter onset and offset responses within bitter-responsive ROIs in anterior, intermediate, and posterior frames (Figure 6D-E). Denatonium onset produced a significantly stronger response than water in each of these ROIs at concentrations as low as 100 μM (p<.05). Thus, the onset response threshold matched that which we observed in GRNs. At a concentration of 1 mM, an offset response to denatonium was observed in area B_A_ (p<.05), but no significant response was observed in the more posterior B_I_ and B_P_ areas. At a concentration of 10 mM, however, the offset response was clearly observed in all ROIs (p<.05). For lobeline, 100 μM evoked onset activity in areas B_A_ and B_P_, and 1 mM was sufficient to drive a significant offset response in all three ROIs (p<.05; Figure S7D).

### GPN Representation of Bitter Offset Extends Beyond the Bitter Onset-Responsive Region

In addition to observing a bitter offset response within the bitter onset-responsive GPN region, we noticed a separate region in which the responses to bitter offset were conspicuously strong. This area was located lateral and slightly ventral to the onset-responsive region, and typically spanned posterior to intermediate planes within the volume. Spatial overlays of bitter offset-responsive regions with those of bitter onset or water suggested that this region, which we call B_L_, did not respond significantly to these stimuli, and thus may be selective for bitter offset (Figure 6D). Indeed, the magnitude of bitter onset and water responses within B_L_ was small, while the magnitude of the offset response was much greater (Figure 6E, bottom row). As in areas B_I_ and B_P_, the threshold for denatonium offset responses in B_L_ was 10 mM (p<.05).

Intriguingly, the spatial pattern of the GPN bitter offset response also appeared to include a region of neuropil resembling the sweet-responsive area S_L_. However, without a sweet stimulus response to provide a benchmark, we could not delineate this region in our bitter concentration experiments. We therefore performed an experiment comparing bitter onset and offset responses to the response to sucrose (Figure 7A). As we previously observed, the offset-responsive region overlapped the onset-responsive region throughout the imaging volume (Figure 7B, top row). However, in anterior planes, the offset-responsive region also overlapped with the sucrose-responsive region (Figure 7B, bottom row). The spatial overlap between sucrose-responsive and bitter offset-responsive areas appeared restricted to S_L_ and did not extend to S_D_ (Figure 7B, bottom row).

**Figure 7.**
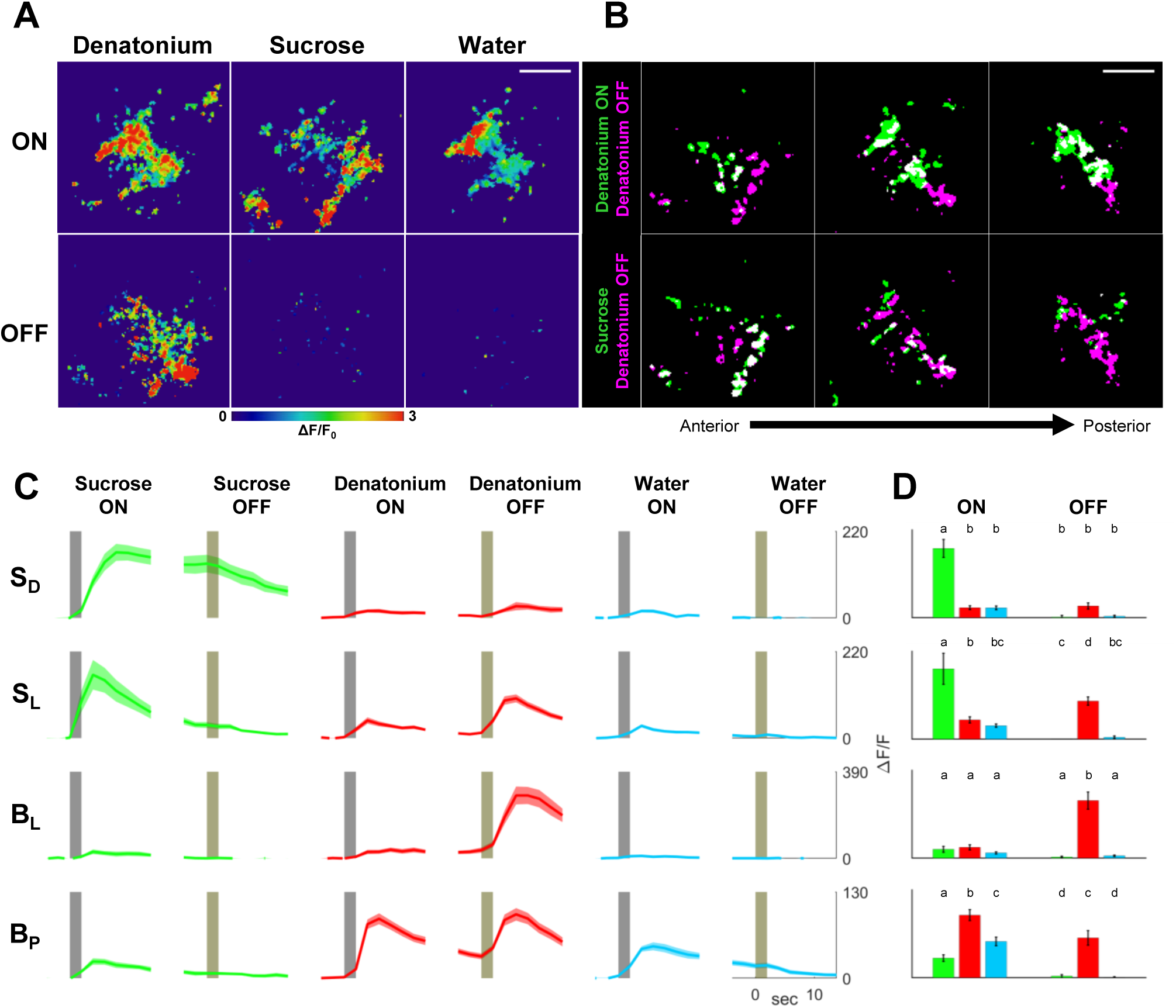
Bitter offset activates a broader set of GPNs than bitter onset. (A) Representative heatmaps showing GPN responses to onset (top row) and offset (bottom row) of denatonium, sucrose, and water. Heatmaps are maximum projections over all planes in the volume; each image represents the response during one trial, and all images are from the same experiment. Heatmaps show ΔF/F value from 0 to 3 as indicated by legend on the right. (B) Representative spatial response maps showing overlap between denatonium offset (magenta) and either denatonium onset (green, top row) or sucrose onset (green, bottom row); overlap is shown in white. (C) GPN responses to onset and offset of sucrose, denatonium, and water within each of four ROIs (n = 10 flies, 3 trials per tastant). (D) Peak responses to onset and offset of each tastant within each ROI. Error bars denote SEM. Letters represent statistically significant differences between groups (p < .05, one-way ANOVA with Tukey post-hoc test). All scalebars 30 μm; scalebar applies to all images in a panel.

To verify this, we compared the magnitude and time course of the responses to sucrose, denatonium, and water within multiple ROIs: the bitter-responsive B_P_, the bitter offset-responsive B_L_, and the sweet-responsive S_L_ and S_D_ (Figure 7C-D). As expected, sucrose drove strong responses in S_L_ and S_D_, with temporal dynamics matching those observed in earlier experiments. While sucrose also elicited activity in areas B_P_ and B_L_, this activity was comparable in magnitude to the water response or weaker than it, and thus not likely to be a response to sweetness. Denatonium onset drove a strong response in area B_P_ and a mild response in area S_L_, but this S_L_ response was comparable to the water response in this region, and thus not likely to reflect a bitter response. Denatonium offset evoked strong activity in both regions B_P_ and B_L_, as previously observed. Further, it elicited a strong response in area S_L_, but only a mild response in area S_D_ (Figure 7C-D). These results suggest that the response to bitter offset involves multiple GPN types, including both bitter onset-responsive GPNs and GPNs specifically tuned to bitter offset, and may potentially include a subset of sweet-responsive GPNs as well.

## DISCUSSION

A central question in taste coding is how the neural representation of sweet and bitter is maintained beyond the sensory neurons at the first order of the circuit. The present study is the first to investigate taste responses selectively in second-order neurons in *Drosophila*, a strategy enabled by a novel configuration of *trans-*Tango. We found evidence that sweet and bitter tastants activate distinct second-order GPNs, just as they do for first-order GRNs. Yet we also found evidence that the representation of bitter in GPNs is somewhat complex. While the GPN response to bitter tastants is strikingly similar to the response to water, bitter tastants are distinguished by their ability to evoke activity upon both onset and offset. The spatial distribution of this offset response is restricted to bitter GRNs in the first order but extends beyond the onset region in the second order, and may recruit GPNs that also respond to sweet. Our results are consistent with a mixed model of taste coding, in which some GPNs may function as labeled lines for particular taste qualities, while others respond to multiple qualities of the stimulus.

Beyond these findings on taste coding, our study demonstrates more generally that *trans-*Tango can fill a niche in analyzing neural circuit function. While a plethora of genetic tools exist for probing circuit function in *Drosophila*, they are limited by the expression patterns of Gal4 driver lines. There is an obvious use in having a line that labels a broad population of neurons at a defined stage of the circuit; for example, the *GH146-Gal4* driver has been extensively used because it labels a broad set of second-order olfactory projection neurons with little background expression (Heimbeck et al., 2001). In many other circuits, though, no comparable driver exists. Using *trans-*Tango initiated from a sparse set of starter neurons, one can selectively image the calcium responses of their downstream projection neurons, as we have done here in the gustatory system. The increasing availability of highly specific split-Gal4 drivers will enable this strategy in a wide variety of systems (Dionne et al., 2018).

Our results suggest that there are GPNs with tuning properties not observed in first-order sensory neurons. For example, some GPNs appear to encode both bitter and water, and thereby integrate signals from different classes of first-order neurons. Further, the water responses of these neurons may be mechanosensory in origin, as PEG elicited similar levels of activation; these GPNs might therefore be integrating across sensory modalities. Other GPNs respond only to bitter offset, which is encoded together with onset by first-order bitter neurons. Thus, some GPNs appear to selectively extract one temporal component of a signal from a single class of first-order neurons. These results suggest that substantial signal convergence and divergence occurs in the transition from the first to the second order of the taste circuit.

We also identified neurons that do not exhibit signatures of such circuit convergence – namely, GPNs selective for sweet taste. Dedicated central pathways for sweet taste may reflect the outsize role sugar has for the animal’s survival. Interestingly, we also found evidence that GPNs represent sweet tastants with both sustained and more transient responses. While it is not immediately apparent what relevance these different temporal patterns may have for behavior, the fact that these two types of responses segregate within the imaging plane suggests that their downstream circuits are distinct as well.

The fact that the fly actively encodes bitter offset raises questions about the relevance of this response for the animal. Distinct onset and offset responses have been observed in other sensory modalities, such as vision (Schiller et al. 1986; Behnia et al., 2014; Strother et al., 2014). In vision, stimulus offset may serve as a cue for the motion of an object, for example. Bitter offset might similarly signal a change in the fly’s gustatory environment. In the wild, flies can encounter a mixture of noxious and nutritious substances at a single feeding site: *Drosophila* feed on microbes that grow on decaying plant matter, and some of these microbes metabolize plant-derived compounds that are normally toxic to flies, and thereby make them safe for the fly’s consumption (Markow & O’Grady 2008). Since many plant toxins are bitter, a fly sampling such a food source would encounter a series of bitter and non-bitter patches; a bitter offset response could thus signal movement off of a toxic substrate and onto a nutritive microbial one. While we do not know the precise number of GPN types involved in the bitter offset response, the distribution of the offset response within areas B_L_, S_L_, and the bitter onset-responsive areas B_A_, B_I_ and B_P_ strongly suggests that several second-order neuron types encode this feature. What function could be served by the distribution of this signal among several channels, rather than keeping it in one, as is the case in the first order? One possibility is that the separation of onset and offset components of the bitter response may endow the fly with greater sensitivity to changes in bitter concentration. In primate vision, separate ON and OFF pathways have been proposed to enable metabolically efficient coding of intensity changes to enhance contrast sensitivity (Schiller et al. 1986) and increase the dynamic range of the system (Kremkow et al., 2014). Likewise, the cockroach has separate olfactory sensory neurons tuned to odor onset and offset, which increase the animal’s ability to detect small concentration changes (Burgstaller and Tichy, 2011). The bitter offset pathway in *Drosophila* might perform a similar function, enabling the fly to efficiently navigate concentration gradients of bitter compounds on heterogeneous food sources. Finally, since this parallel ON/OFF pathway circuit motif has been observed in several sensory systems, including vision (Schiller et al., 1986; Strother et al., 2014), olfaction (Chalasani et al., 2007; Burgstaller and Tichy, 2011), touch (Iggo and Ogawa, 1977), and thermosensation (Gallio et al., 2011), it may have general utility in sensory coding.

Another possibility is that bitter onset and offset may have opposing hedonic valences to the fly. The cessation of punishment is known to act as a reward in conditioning experiments (Gerber et al., 2014). Because bitter taste is aversive, bitter offset may thus be interpreted as rewarding. Separating these components of the bitter taste response may allow for differential input to dopaminergic neurons of the mushroom body that mediate aversive and appetitive conditioning. The intriguing observation that the bitter offset response overlaps with the sweet response in the area we call S_L_ is consistent with this idea. If there is a GPN that underlies both the bitter offset and sweet responses in this area, this neuron may functionally encode the valence of the stimulus. Yet even if these responses are caused by separate sweet and bitter offset GPNs, their overlapping projections in this region may indicate that these GPNs converge onto common third-order neurons in the circuit.

Together, our results revealed a hybrid model of taste quality coding in the *Drosophila* gustatory system. The GPN population we studied includes both labeled line and convergent representations of taste, and distributes information about stimulus timing across multiple neural pathways. It remains to be seen how this manner of encoding gustatory information is used by downstream circuits to sculpt the behavior of the fly.

## ACKNOWLEDGMENTS

This work was supported by NIH grants R01DC017146 and R01MH105368 as well as by internal funds from Brown University: Carney Innovation Award and OVPR Seed Award (GB). NJS was supported by grant number DGE1058262 from the National Science Foundation Graduate Research Fellowship Program. We thank Alexander Fleischmann and members of the Barnea lab for critical reading of the manuscript.

## AUTHOR CONTRIBUTIONS

N.J.S. and G.B. designed the experiments; N.J.S., J.D.F., G.G.H. and M.T. performed the experiments; N.J.S., J.D.F., G.G.H., M.T. and G.B. analyzed data; N.J.S., and G.B. wrote the manuscript. J.D.F. and M.T. commented on the manuscript.

## STAR METHODS

### RESOURCE AVAILABILITY

#### Lead Contact

Further information and requests for resources and reagents should be directed to and will be fulfilled by the Lead Contact, Gilad Barnea.

#### Materials Availability

Fly lines generated in this study are available upon request.

#### Data and Code Availability

Data and code generated in this study are available upon request.

## EXPERIMENTAL MODEL AND SUBJECT DETAILS

### Fly strains

*Drosophila melanogaster* lines used in this study were raised on standard cornmeal-agar media (with tegosept anti-fungal agent). Flies were maintained in humidity-controlled chambers kept at 18°C and set to a 12-hour light/dark cycle. The following fly lines were used: *Gr64f-Gal4* (Dahanukar et al., 2007), *Gr66a-Gal4* (Scott et al., 2001), *trans-Tango* (Talay et al., 2017), *UAS-myrGFP, QUAS-mtdTomato(3xHA)* (Talay et al., 2017), *20xUAS-IVS-jGCaMP7b* (Dana et al., 2019), *QUAS-GCaMP6s* (this study).

### Genotypes

The genotypes used in each figure are as follows:

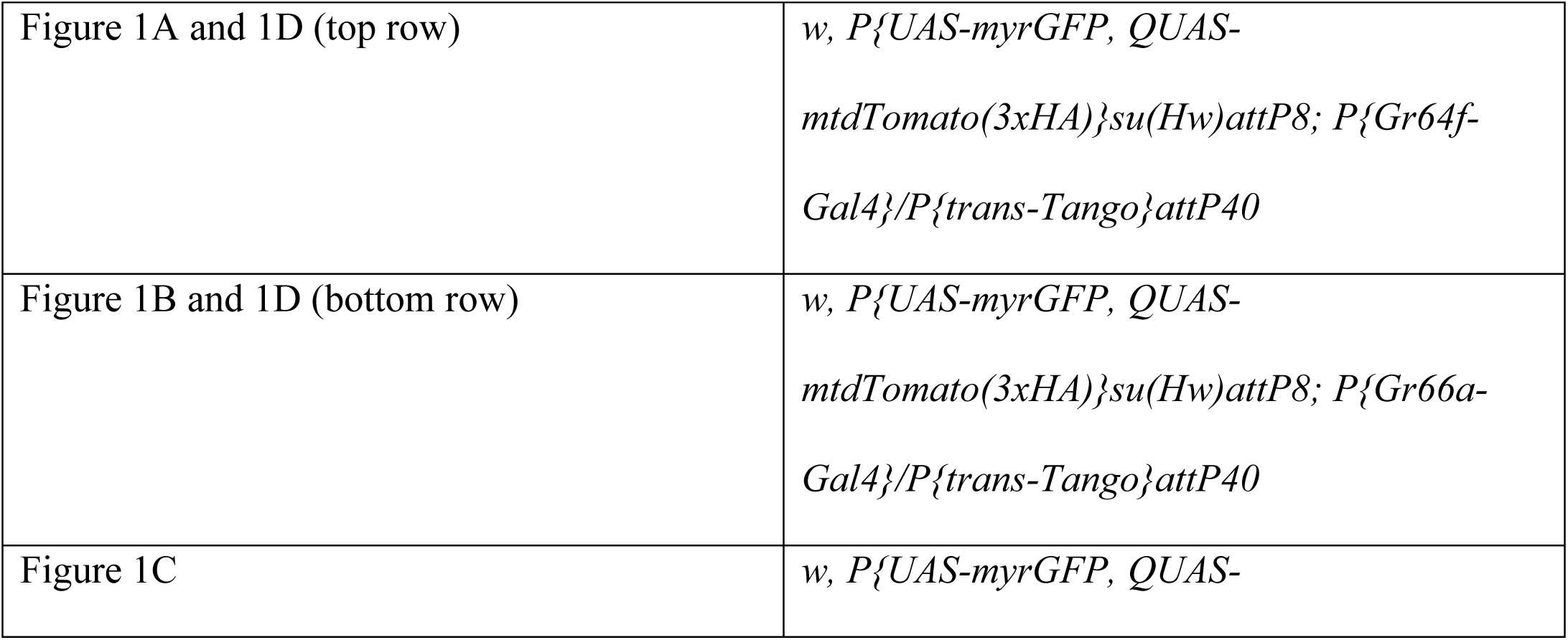
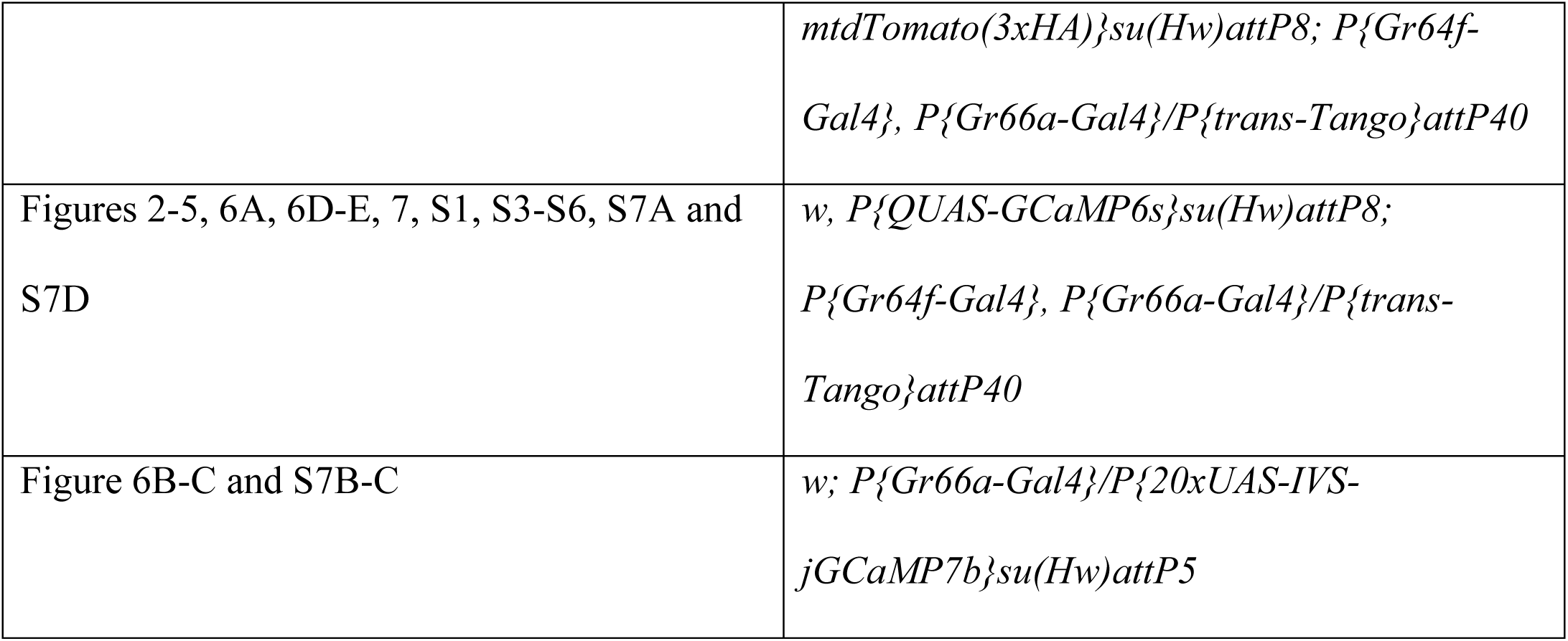

### Generation of transgenic fly strains

The QUAS-GCaMP6s construct was designed with several elements included to boost reporter expression levels (Pfeiffer et al., 2010; Pfeiffer et al., 2012): a 10xQUAS site, 5’UTR intervening sequence (IVS), 5’UTR syn21 sequence, codon optimized GCaMP6s, and a p10 terminator. The plasmid used to generate reporter flies was constructed by PCR, restriction digest and Gibson Assembly. The IVS, syn21 and p10 sequences, obtained from pJFRC81 (Addgene, 36432), and 10xQUAS site and GCaMP6s codon optimized sequences, obtained by synthesis (GeneArt, Thermo Fisher; seed sequence: (Chen et al., 2013)), were amplified with the appropriate Gibson overlaps. The plasmid backbone was obtained by digesting UAS-myrGFP-QUAS-mtdTomato(3xHA) (Talay et al, 2017) to replace the UAS-myrGFP and mtdTomato(3xHA) components. These fragments were assembled by Gibson assembly (New England Biolabs) cloned and sequence verified before injection.

## METHOD DETAILS

### Immunohistochemistry

Immunohistochemistry was performed as previously described (Talay et al., 2017). Images are of brains of adult males aged 15-20 days. The following primary antibodies were used: rabbit anti-GFP (Thermo Fisher Scientific, A11122; 1:1,000), rat anti-HA (Roche, 11867423001; 1:100), mouse anti-Brp (nc82; DSHB; 1:50). The following secondary antibodies were used, all at 1:1000 dilution: donkey anti-rabbit (Alexa Fluor 488), goat anti-rat (Alexa Fluor 555), donkey anti-mouse (Alexa Fluor 647). Imaging was performed with a Zeiss confocal microscope (LSM800) using ZEN software (Zeiss, version 2.1). Images were further formatted using Fiji software (http://fiji.sc).

### Calcium imaging

All imaging was performed on adult male flies. All flies were aged 10-15 days to maximize *trans*-Tango-dependent GCaMP signal. In starvation experiments, flies were wet-starved in a vial with a wet Kimwipe for 22-26 hours before imaging.

Preparation of flies for calcium imaging was based on previously described methods (Yapici et al., 2016). An imaging chamber was created by drilling a hole into a small plastic tissue culture dish. A piece of clear packing tape was used to cover the hole so that the non-adhesive side formed the basin of the dish. Flies were anesthetized with CO_2_ and placed onto the sticky side of the tape. A human hair placed across the neck secured the fly in place; the hair was held in place using thin strips of tape. Flies extend their proboscis as they recover from CO_2_ anesthesia, and during this process the proboscis was glued in an extended position using UV-curing glue (Loctite). Forelegs were removed along with tarsi of other legs to prevent both interference with the stimulus and accidental activation of tarsal gustatory neurons. A window was cut into the tape using a syringe needle, and the head capsule was inserted through the window and glued into place with UV-curing glue. The dish was filled with artificial hemolymph-like (AHL) solution (108 mM NaCl, 5 mM KCl, 2 mM CaCl_2_, 8.2 mM MgCl_2_, 4 mM NaHCO_3_, 1mM NaH_2_PO_4_, 15 mM ribose, 5 mM HEPES, pH adjusted to 7.2-7.5, osmolarity adjusted to ∼275 mOsm). Ribose was substituted for other sugars in the AHL to avoid spurious activation of sugar-sensing neurons (Marella et al., 2006). A small window was cut into the fly cuticle to expose the imaging area, and obstructing tissue and air sacs were removed. Antennae and muscle 16 were removed to reduce motion. When imaging the SEZ, the esophagus was cut to reduce motion and expose the imaging area.

Two-photon calcium imaging was performed on a Scientifica multiphoton galvo system. The excitation wavelength used was 920 nm, and laser power at the objective was kept between 15-20 mW. Imaging data was collected using SciScan software (Scientifica). A water-immersion objective (Nikon, 16x, 0.8 NA) was used and imaging volumes consisting of 12 planes of depth with 8-9 μm spacing between planes were acquired with 256×256 pixel resolution per plane, using 391 nm pixel size for GPN imaging and 521 nm pixel size for GRN imaging. Volumes were collected at a ∼0.5 Hz volume rate (∼6 Hz frame rate).

Each imaging session consisted of three trials per tastant, where each trial consisted of one stimulus delivery, held for several seconds. For experiments in which stimulus offset was monitored, the stimulus hold time was varied between 20, 25, and 30 seconds; for all other experiments, hold time was 8-10 seconds. To ensure the fly remained responsive throughout the imaging session, overall imaging time per fly was kept under 2 hours. Trials using the same tastant were therefore performed consecutively to reduce overall imaging time. Stimulus presentations were spaced out by ∼2 minutes to minimize habituation.

Tastant solutions were delivered to the proboscis during imaging using a Drummond Nanoject II microinjector mounted on a micromanipulator (Scientifica). In each trial, stimuli were manually delivered after collecting at least 10 baseline imaging volumes. Tastant delivery to the proboscis was monitored using a Point Grey Firefly camera equipped with an Infinistix lens and a shortpass IR filter (850nm OD 4.0 shortpass filter, Edmund Optics). An LED flash was triggered by the software upon the start of an imaging trial, and the frame of tastant delivery was calculated for each trial based on elapsed time since the flash. Tastant solutions used for imaging were used at the following concentrations: polyethylene glycol (3,350 g/mol) (20% w/v); fructose, glucose, maltose, sucrose, trehalose (1 M); caffeine, papaverine hydrochloride, quinine hydrochloride dihydrate (10 mM); denatonium benzoate and lobeline hydrochloride were used at 10 μM, 100 μM, 1 mM, and 10 mM for concentration experiments and at 1 mM for bitter panel experiments; denatonium was used at 10 mM in experiments in Figure 2 and Figure 7.

## QUANTIFICATION AND STATISTICAL ANALYSIS

### Calcium imaging analysis

Raw images were preprocessed using Fiji functions “Subtract Background” (rolling ball radius of 50 pixels) and “Gaussian Blur” (radius of 2 standard deviations). In MATLAB, motion correction was performed on each plane in the imaging volume using the NoRMCorre algorithm (Pnevmatikakis & Giovannucci, 2017).

We performed imaging analysis in MATLAB, using a strategy inspired by a previously described method (Liang et al., 2013). We limited our analysis to a general ROI within the imaging volume consisting only of responsive pixels. We determined this ROI in three steps. First, for each trial we determined which pixels in the imaging volume showed a significant response to the stimulus. For each pixel, we defined a significant response as three standard deviations above its baseline fluorescence level. We considered the three imaging volumes following stimulus delivery to comprise the response to stimulus onset, and used the mean of ten volumes before stimulus as the baseline. To reduce noise, we considered only those pixels that showed significant responses in all three volumes following stimulus delivery. Second, we defined specific ROIs for each tastant, representing the average region activated by that tastant across trials. We considered only those pixels showing significant responses in two out of three trials for that tastant, then median filtered the resulting image and removed from consideration any border pixels that did not remain in frame for all trials in the experiment. Third, we created the general ROI consisting of all responsive pixels in an experiment by taking the union of all tastant-specific ROIs.

To determine peak responses for each trial, we took the three post-stimulus volumes and considered only those pixels within the general ROI, calculated the maximum ΔF/F across these volumes for each pixel showing a significant response, then median filtered the resulting image. To display this peak response as a heatmap, we exported the image to Fiji, created a maximum Z projection, and used the “physics” LUT with a max of 300% ΔF/F. To calculate correlations between trials, we took the peak responses for each trial, converted ΔF/F values of all pixels within the general ROI to vectors, and computed Pearson’s correlation coefficient between these vectors. This produced correlation values for every trial pair for each fly; the correlation matrices displayed in each figure depicts the mean of these inter-trial correlation values across all flies. To generate maps of spatial patterns of tastant responses, we displayed all pixels that were active in any of the three trials for a given tastant. To define ROIs for calculating fluorescence traces and response magnitude, we took motion-corrected single imaging planes and used the maps of spatial patterns of responses as a guide to manually draw boundaries around regions that exhibited significant responses to a tastant.

For trials where we analyzed both stimulus onset and offset, the method for defining the general ROI was slightly modified. Stimulus onset-responsive regions and ROIs were calculated the same way. In addition to determining stimulus onset-responsive pixels, we also determined offset-responsive pixels. The analysis period for defining offset-responsive pixels was the volume when stimulus removal occurred plus the following three volumes; significant responses were considered to be those exhibiting activity greater than three standard deviation above baseline level. The offset baseline level was defined as the mean pixel fluorescence of the five volumes prior to stimulus removal. In the onset/offset experiments, we generated ROIs for each tastant-on/off pairing (e.g. separate ROIs for denatonium-on and denatonium-off), and considered the union of all tastant-on/off ROIs as the general ROI for that experiment. For first-order experiments, to reduce background noise we only used tastant-on ROIs to construct the general ROI; there is no apparent subregion of GRN axons that is selectively offset-tuned, so this modification did not exclude any responsive areas.

### Statistics

To determine whether correlations between responses to two tastants were statistically significant, we computed Pearson’s r between each pair of trials for those tastants; since we ran three trials for each tastant during an imaging session, for every pair of tastants there were nine pairs of trials, and we took the mean of these nine Pearson’s r values to determine a correlation value for that tastant pair for each fly. We ran a one sample t-test on these correlation values against a mean of 0 for each tastant pair and used a Bonferroni correction to account for multiple comparisons. To determine whether differences between taste responses within an ROI were statistically significant, we performed one-way ANOVA tests with Tukey post-hoc tests to account for multiple comparisons. To determine significant concentration threshold levels for bitter onset and offset responses, we performed two-sample one-tailed t-tests with Bonferroni correction, where the experimental hypothesis was that the bitter response was greater than the water response within the ROI. All statistics were performed in MATLAB.

---

**Figure S1.**
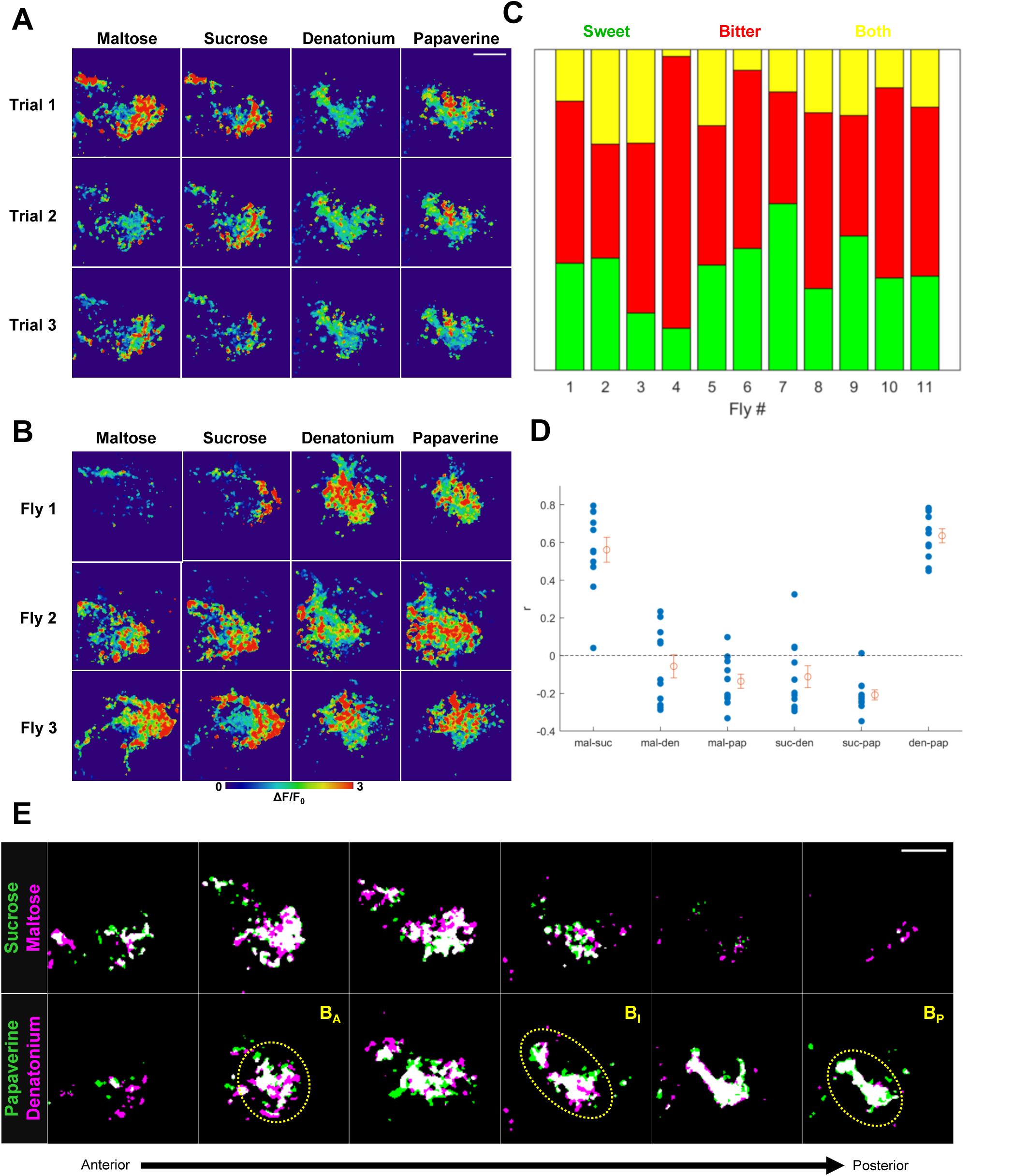
Consistency of GPN responses between trials and tastants, related to Figure 2. (A) Heatmaps of peak responses to a tastant during each of three trials for an example fly. Top row is the first trial, middle row is second, bottom row is third. Heatmaps are maximum projections over all planes in the volume. Heatmaps show ΔF/F value from 0 to 3 as indicated by legend. (B) Heatmaps for three different flies. (C) Proportion of pixels responding to each taste quality. Green: proportion of pixels only responding to sweet stimuli; red: proportion only responding to bitter; yellow: proportion responding to both sweet and bitter. Each column shows results for one fly. (D) Inter-tastant correlations for individual flies. Each blue dot represents average correlation between trials of a given tastant pair for a single fly. Orange dot denotes mean; error bars denote SEM. Sucrose/maltose and denatonium/papaverine correlations are significantly greater than zero (p < .05, one sample t-test, Bonferroni correction). (E) Overlaid spatial response maps for tastants of the same quality. Top row: sucrose (green) and maltose (magenta), bottom row: papaverine (green) and denatonium (magenta); overlap shown in white. ROIs for areas B_A_, B_I_, and B_P_ shown on bottom row. All scalebars 30 μm; scalebar in A applies to A and B, and scalebar in D applies to all images in the panel.

**Figure S2.**
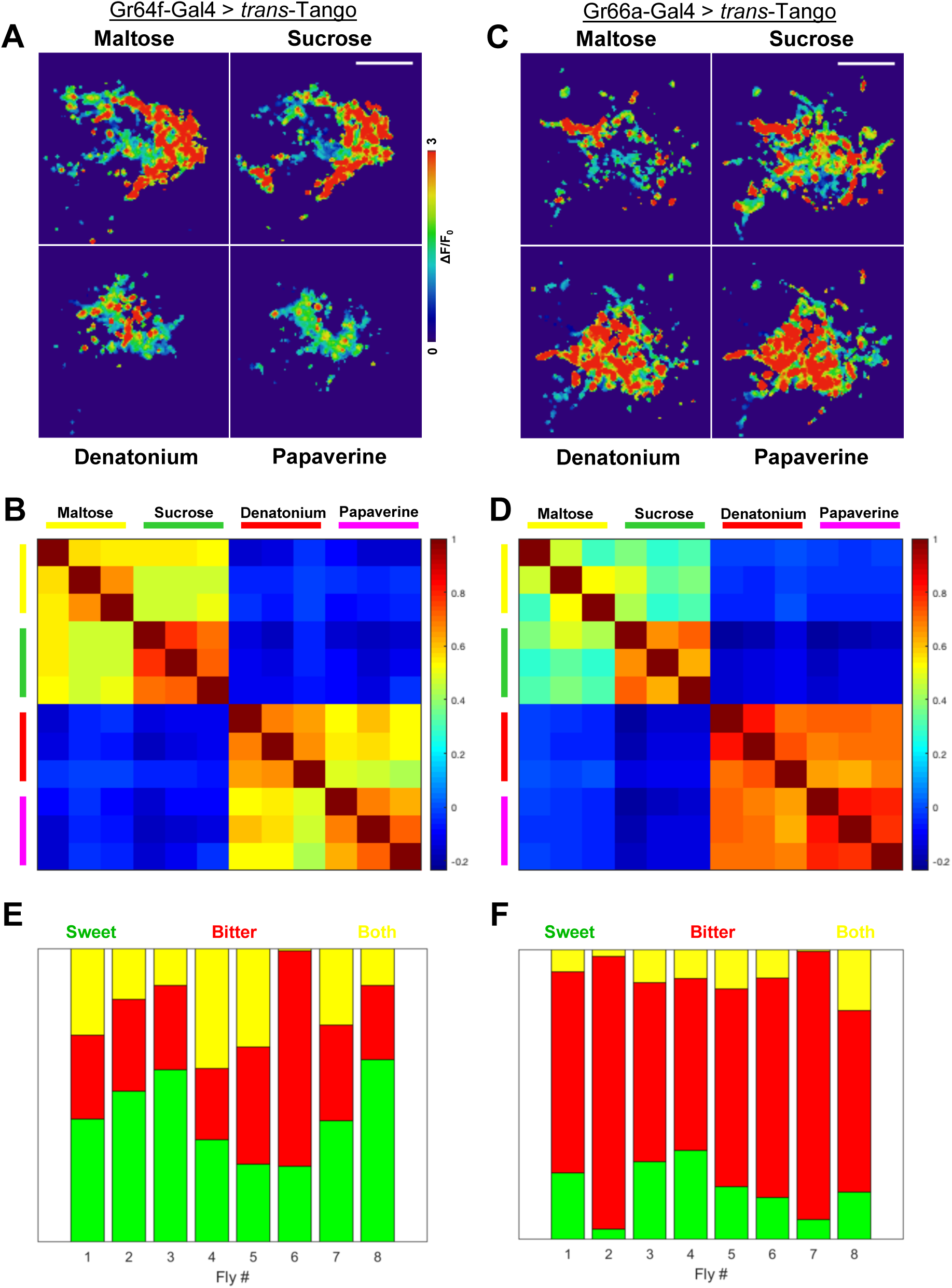
Taste responses of GPNs labeled by *trans*-Tango driven by either *Gr64f-Gal4* or *Gr66a-Gal4*, related to Figure 2. (A) Representative heatmaps showing responses of GPNs labeled by *Gr64f-Gal4* > *trans-*Tango to sweet (maltose, sucrose) and bitter (denatonium, papaverine) tastants. (B) Correlation matrix showing pairwise inter-trial correlations of peak responses of GPNs labeled by *Gr64f-Gal4* > *trans*-Tango; group-averaged (n = 8 flies). (C) Heatmaps as in A, but for GPNs labeled by *Gr66a-Gal4* > *trans*-Tango. (D) Correlation matrix as in B, but for GPNs labeled by *Gr64f-Gal4* > *trans*-Tango (n = 8 flies). (E) Proportion of pixels responding to each taste quality, for *Gr64f-Gal4* > *trans*-Tango. Green: proportion of pixels only responding to sweet stimuli; red: proportion only responding to bitter; yellow: proportion responding to both sweet and bitter. Each column shows results for one fly. (F) Proportion of pixels responding to each taste quality, as in E, for *Gr66a-Gal4* > *trans*-Tango. All scalebars 30 μm; scalebar applies to all images in a panel. Colors same as in E.

**Figure S3.**
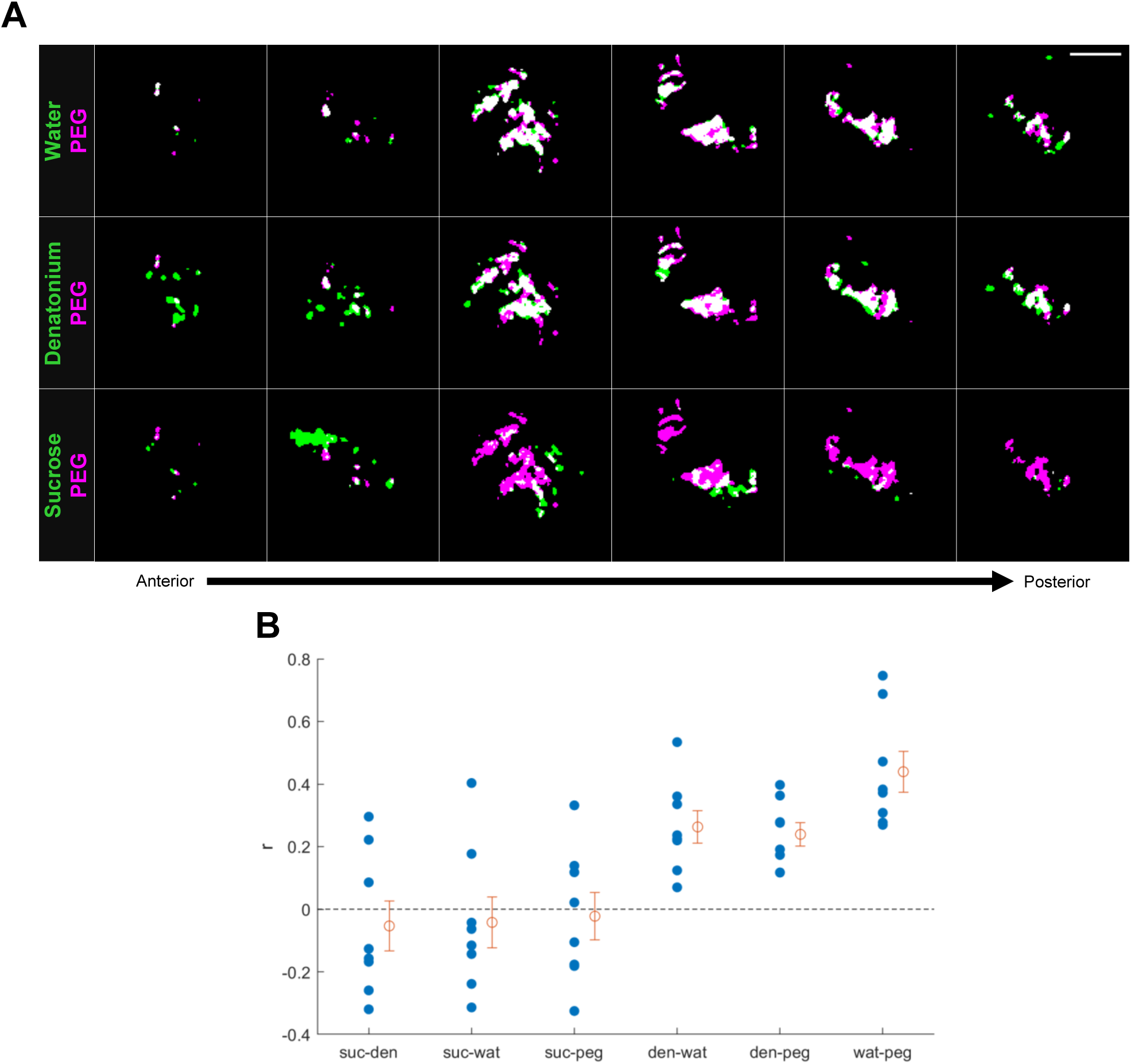
PEG and water activate similar GPNs, related to Figure 3. (A) Representative overlaid spatial maps of PEG-evoked responses (magenta) with those of water (green, top row), denatonium (green, middle row), and sucrose (green, bottom row). (B) Inter-tastant correlations for individual flies. Each blue dot represents average correlation between trials of a given tastant pair for a single fly. Orange dot denotes mean; error bars denote SEM. Denatonium/water, denatonium/PEG, and water/PEG correlations are significantly greater than zero (p < .05, one sample t-test, Bonferroni correction). Scalebar 30 μm and applies to all panels.

**Figure S4.**
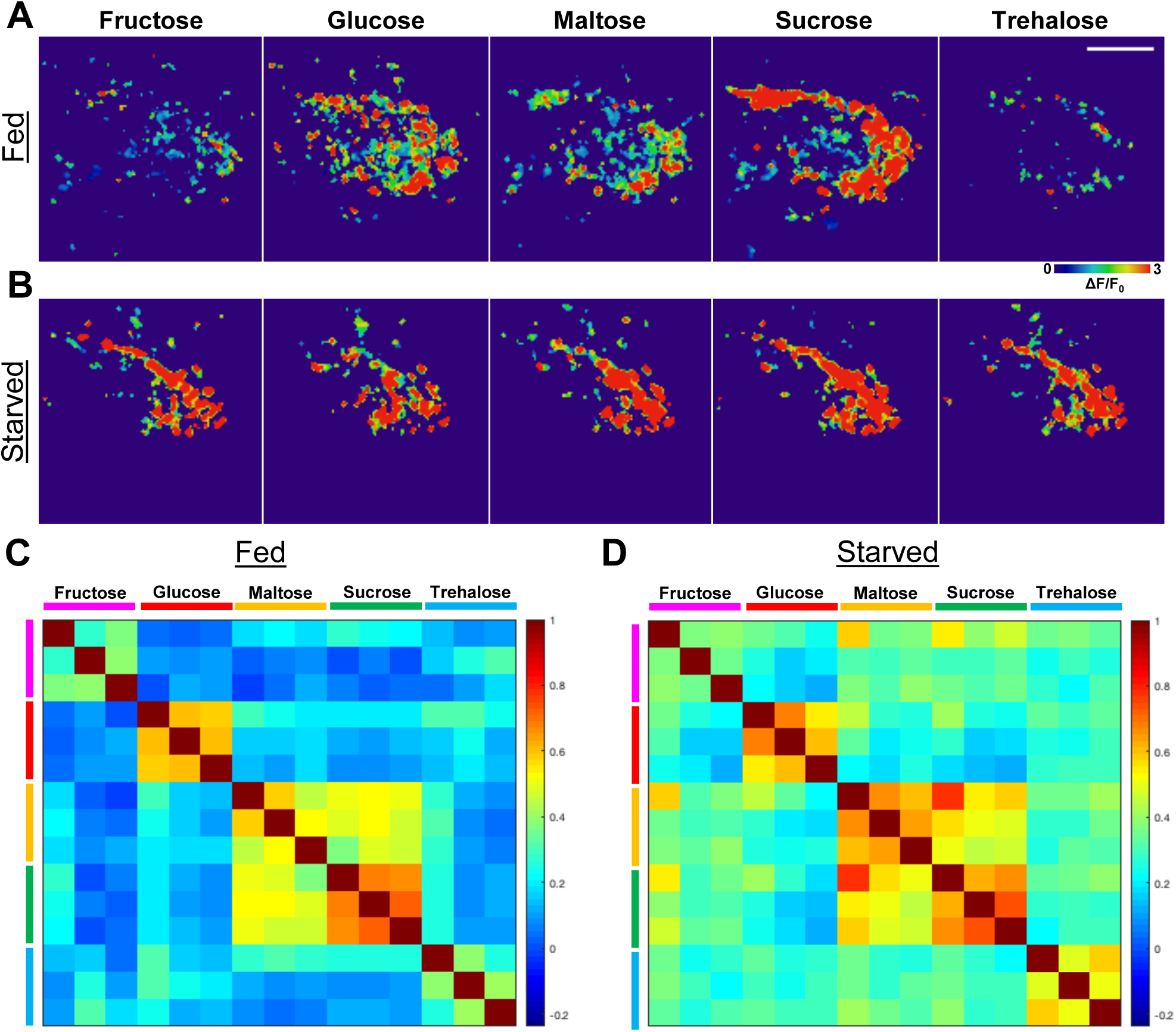
Responses to sweet tastants excluding water-responsive pixels, related to Figure 4. (A) Heatmaps of same responses to sweet tastants shown in Figure 4A, but excluding the water-responsive pixels. (B) Heatmaps of same responses to sweet tastants shown in Figure 4B, but excluding the water-responsive pixels. (C) Correlation matrix of GPN responses to sweet tastants in fed flies, as shown in Figure 4C, but limited to region excluding water-responsive pixels. (D) Correlation matrix of GPN responses to sweet tastants in starved flies, as shown in Figure 4D, but limited to region excluding water-responsive pixels. Scalebar 30 μm and applies to all panels in figure.

**Figure S5.**
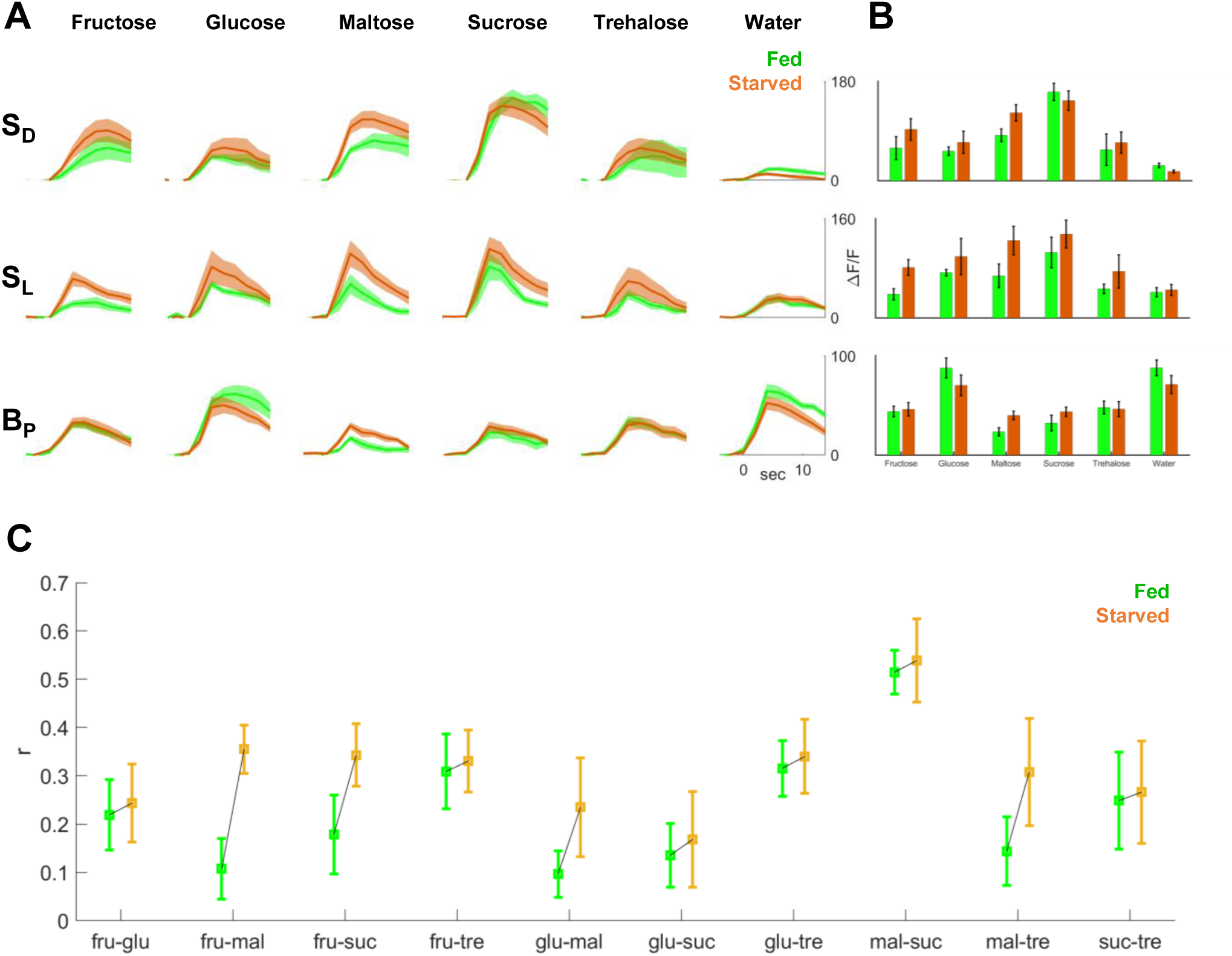
Effects of starvation on GPN responses to sweet tastants, related to Figure 4. (A) Reponses to sweet tastants in the two sweet-responsive regions, S_D_ and S_L_, and water/bitter-responsive region B_P_. Green traces indicate responses for fed flies, orange traces indicate responses for starved flies (n = 8 fed; n = 8 starved). (B) Max ΔF/F for each tastant in each ROI. (C) Inter-tastant correlations are equal or greater for starved flies than fed flies. Squares denote mean correlation across individuals; error bars denote SEM. Green: values for fed flies, orange: values for starved flies.

**Figure S6.**
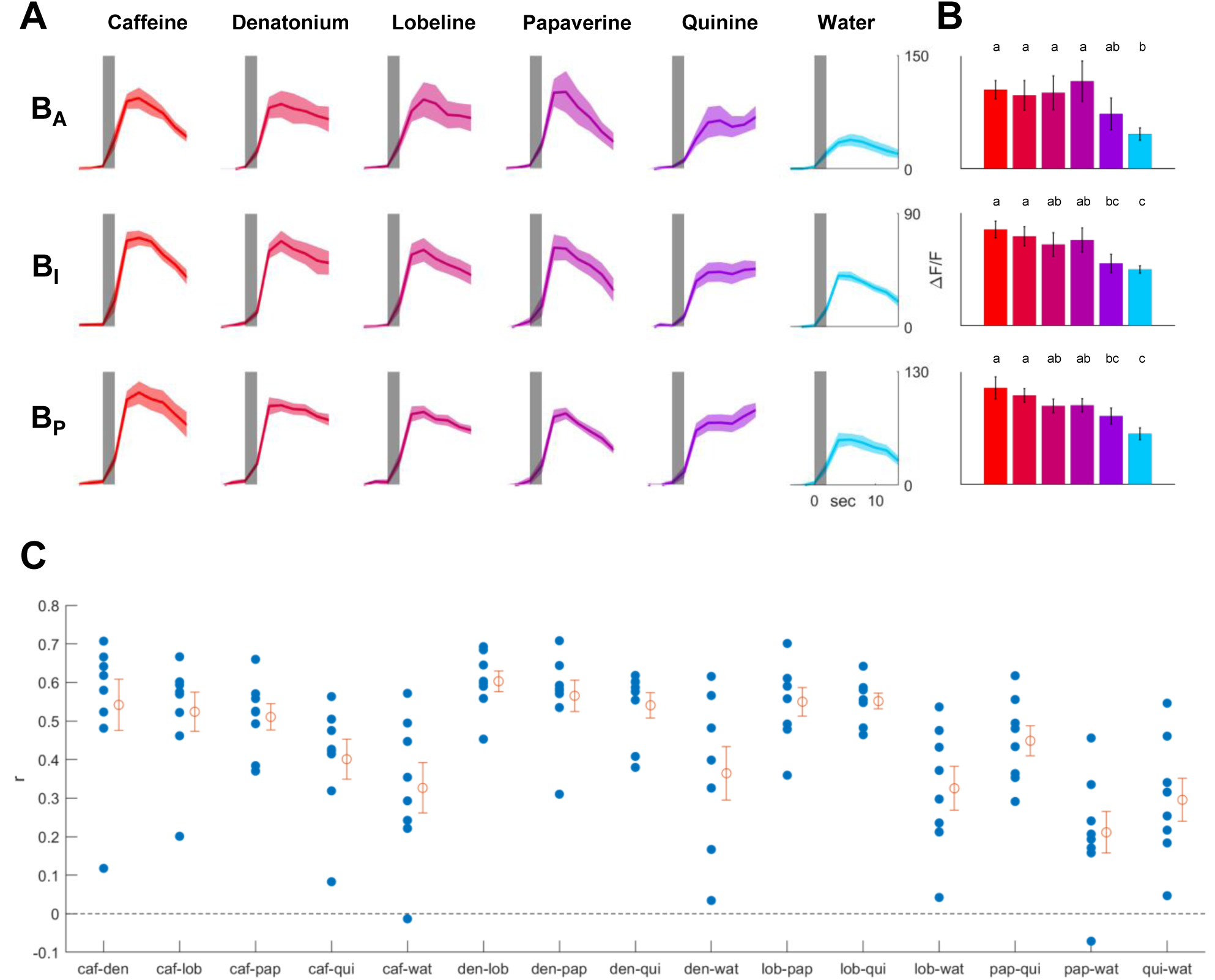
Responses to bitter tastants within each bitter ROI, related to Figure 5. (A) Reponses to bitter tastants in each of three bitter ROIs (n = 8). (B) Max ΔF/F for each tastant in each ROI. Letters represent statistically significant differences between groups (p < .05, one-way ANOVA with Tukey post-hoc test). (C) Inter-tastant correlations for individual flies. Each blue dot represents average correlation between trials of a given tastant pair for a single fly. Orange dot denotes mean; error bars denote SEM. Except for papaverine/water, correlations for all pairs were significantly greater than zero (p < .05, one sample t-test compared to 0, Bonferroni correction).

**Figure S7.**
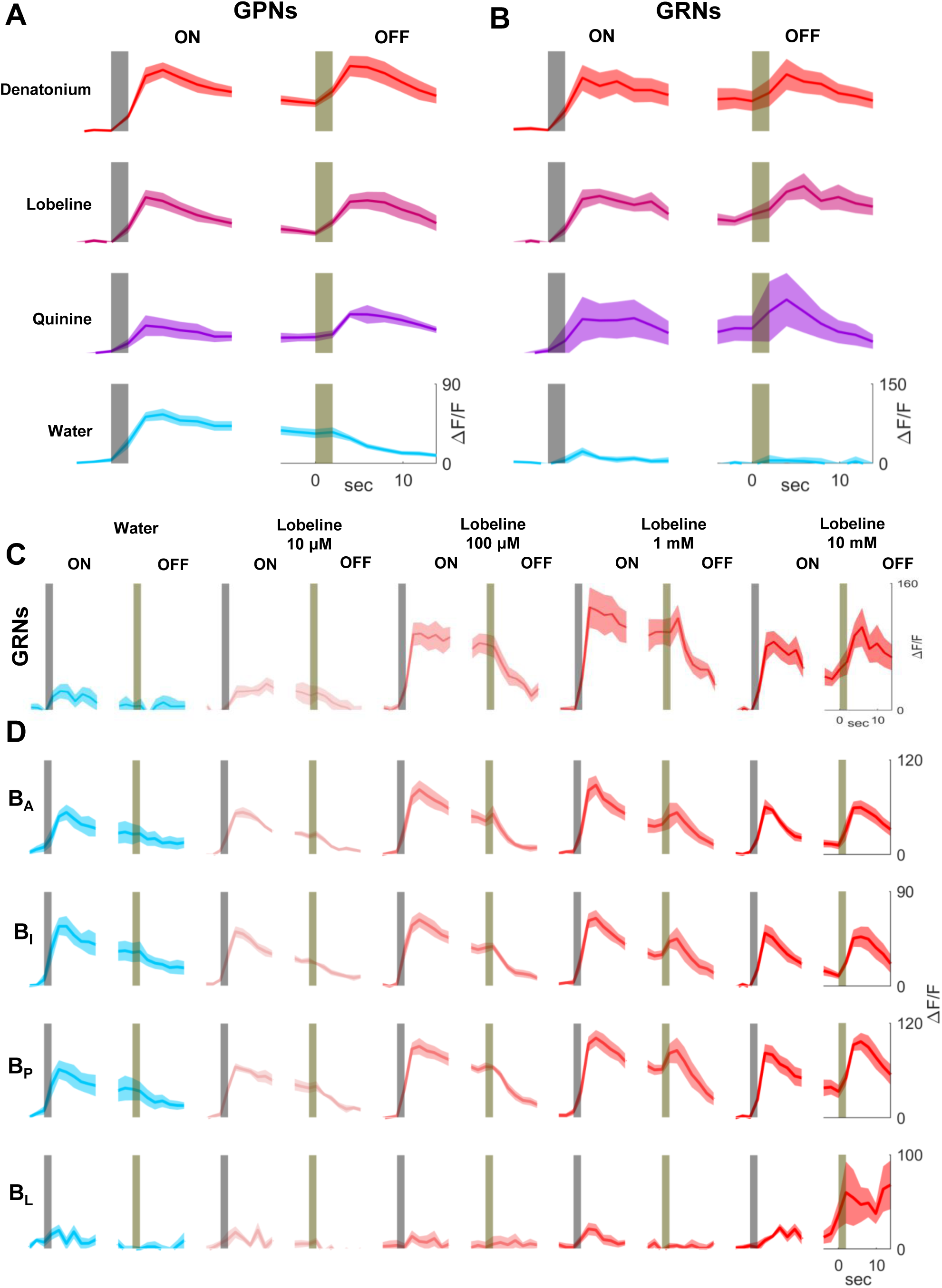
Bitter offset response to various bitter tastants, related to Figure 6. (A) GPN responses to onset and offset of bitter compounds and water (denatonium: n = 6; lobeline: n = 6; quinine: n = 3, water: n = 6). (B) GRN responses to onset and offset of bitter compounds and water (denatonium: n = 5; lobeline: n = 6; quinine: n = 2, water: n = 5). (C) GRN responses to increasing concentrations of lobeline (n = 6). (D) GPN responses to increasing concentrations of lobeline in different ROIs (n = 6).

## REFERENCES

Behnia, R., Clark, D.A., Carter, A.G., Clandinin, T.R., and Desplan, C. (2014). Processing properties of on and off pathways for Drosophila motion detection. Nature 512, 427–430.

Bohra, A.A., Kallman, B.R., Reichert, H., and VijayRaghavan, K. (2018). Identification of a Single Pair of Interneurons for Bitter Taste Processing in the Drosophila Brain. Current Biology 28, 847-858.e3.

Burgstaller, M., and Tichy, H. (2011). Functional asymmetries in cockroach ON and OFF olfactory receptor neurons. Journal of Neurophysiology 105, 834–845.

Cameron, P., Hiroi, M., Ngai, J., and Scott, K. (2010). The molecular basis for water taste in Drosophila. Nature 465, 91–95.

Chalasani, S.H., Chronis, N., Tsunozaki, M., Gray, J.M., Ramot, D., Goodman, M.B., and Bargmann, C.I. (2007). Dissecting a circuit for olfactory behaviour in Caenorhabditis elegans. Nature 450, 63–70.

Chen, T.W., Wardill, T.J., Sun, Y., Pulver, S.R., Renninger, S.L., Baohan, A., Schreiter, E.R., Kerr, R.A., Orger, M.B., Jayaraman, V., et al. (2013). Ultrasensitive fluorescent proteins for imaging neuronal activity. Nature 499, 295–300.

Clyne, P.J., Warr, C.G., and Carlson, J.R. (2000). Candidate taste receptors in Drosophila. Science 287, 1830–1834.

Dahanukar, A., Foster, K., Van der Goes van Naters, W.M., and Carlson, J.R. (2001). A Gr receptor is required for response to the sugar trehalose in taste neurons of Drosophila. Nature Neuroscience 4, 1182–1186.

Dahanukar, A., Lei, Y.T., Kwon, J.Y., and Carlson, J.R. (2007). Two Gr Genes Underlie Sugar Reception in Drosophila. Neuron 56, 503–516.

Dana, H., Sun, Y., Mohar, B., Hulse, B.K., Kerlin, A.M., Hasseman, J.P., Tsegaye, G., Tsang, A., Wong, A., Patel, R., et al. (2019). High-performance calcium sensors for imaging activity in neuronal populations and microcompartments. Nature Methods 16, 649–657.

Dionne, H., Hibbard, K.L., Cavallaro, A., Kao, J.C., and Rubin, G.M. (2018). Genetic reagents for making split-GAL4 lines in Drosophila. Genetics 209, 31–35.

Dunipace, L., Meister, S., McNealy, C., and Amrein, H. (2001). Spatially restricted expression of candidate taste receptors in the Drosophila gustatory system. Current Biology 11, 822–835.

Gallio, M., Ofstad, T.A., Macpherson, L.J., Wang, J.W., and Zuker, C.S. (2011). The coding of temperature in the Drosophila brain. Cell 144, 614–624.

Gerber, B., Yarali, A., Diegelmann, S., Wotjak, C.T., Pauli, P., and Fendt, M. (2014). Pain-relief learning in flies, rats, and man: Basic research and applied perspectives. Learning and Memory 21, 232–252.

Harris, D.T., Kallman, B.R., Mullaney, B.C., and Scott, K. (2015). Representations of Taste Modality in the Drosophila Brain. Neuron 86, 1449–1460.

Heimbeck, G., Bugnon, V., Gendre, N., Keller, A., and Stocker, R.F. (2001). A central neural circuit for experience-independent olfactory and courtship behavior in Drosophila melanogaster. Proceedings of the National Academy of Sciences of the United States of America 98, 15336–15341.

Iggo, A., and Ogawa, H. (1977). Correlative physiological and morphological studies of rapidly adapting mechanoreceptors in cat’s glabrous skin. The Journal of Physiology 266, 275–296.

Inagaki, H.K., Ben-Tabou De-Leon, S., Wong, A.M., Jagadish, S., Ishimoto, H., Barnea, G., Kitamoto, T., Axel, R., and Anderson, D.J. (2012). Visualizing neuromodulation in vivo: TANGO-mapping of dopamine signaling reveals appetite control of sugar sensing. Cell 148, 583–595.

Kain, P., and Dahanukar, A. (2015). Secondary Taste Neurons that Convey Sweet Taste and Starvation in the Drosophila Brain. Neuron 85, 819–832.

Kim, H., Kirkhart, C., and Scott, K. (2017). Long-range projection neurons in the taste circuit of Drosophila. ELife 6.

Kremkow, J., Jin, J., Komban, S.J., Wang, Y., Lashgari, R., Li, X., Jansen, M., Zaidi, Q., and Alonso, J.M. (2014). Neuronal nonlinearity explains greater visual spatial resolution for darks than lights. Proceedings of the National Academy of Sciences of the United States of America 111, 3170–3175.

Kvello, P., Jørgensen, K., and Mustaparta, H. (2010). Central gustatory neurons integrate taste quality information from four appendages in the moth Heliothis virescens. Journal of Neurophysiology 103, 2965–2981.

Liang, L., Li, Y., Potter, C.J., Yizhar, O., Deisseroth, K., Tsien, R.W., and Luo, L. (2013). GABAergic Projection Neurons Route Selective Olfactory Inputs to Specific Higher-Order Neurons. Neuron 79, 917–931.

Liman, E.R., Zhang, Y.V., and Montell, C. (2014). Peripheral coding of taste. Neuron 81, 984–1000.

Marella, S., Fischler, W., Kong, P., Asgarian, S., Rueckert, E., and Scott, K. (2006). Imaging taste responses in the fly brain reveals a functional map of taste category and behavior. Neuron 49, 285–295.

Markow, T.A., and O’Grady, P. (2008). Reproductive ecology of Drosophila. Functional Ecology 22, 747–759.

Miyazaki, T., Lin, T.Y., Ito, K., Lee, C.H., and Stopfer, M. (2015). A gustatory second-order neuron that connects sucrose-sensitive primary neurons and a distinct region of the gnathal ganglion in the Drosophila brain. Journal of Neurogenetics 29, 144–155.

Moon, S.J., Köttgen, M., Jiao, Y., Xu, H., and Montell, C. (2006). A Taste Receptor Required for the Caffeine Response In Vivo. Current Biology 16, 1812–1817.

Moreira, J.M., Itskov, P.M., Goldschmidt, D., Baltazar, C., Steck, K., Tastekin, I., Walker, S.J., and Ribeiro, C. (2019). Optopad, a closed-loop optogenetics system to study the circuit basis of feeding behaviors. ELife 8.

Musso, P.Y., Junca, P., Jelen, M., Feldman-Kiss, D., Zhang, H., Chan, R.C.W., and Gordon, M.D. (2019). Closed-loop optogenetic activation of peripheral or central neurons modulates feeding in freely moving Drosophila. ELife 8.

Pfeiffer, B.D., Ngo, T.T.B., Hibbard, K.L., Murphy, C., Jenett, A., Truman, J.W., and Rubin, G.M. (2010). Refinement of tools for targeted gene expression in Drosophila. Genetics 186, 735–755.

Pfeiffer, B.D., Truman, J.W., and Rubin, G.M. (2012). Using translational enhancers to increase transgene expression in Drosophila. Proceedings of the National Academy of Sciences of the United States of America 109, 6626–6631.

Pnevmatikakis, E.A., and Giovannucci, A. (2017). NoRMCorre: An online algorithm for piecewise rigid motion correction of calcium imaging data. Journal of Neuroscience Methods 291, 83–94.

Reiter, S., Rodriguez, C.C., Sun, K., and Stopfer, M. (2015). Spatiotemporal coding of individual chemicals by the gustatory system. Journal of Neuroscience 35, 12309–12321.

Schiller, P.H., Sandell, J.H., and Maunsell, J.H.R. (1986). Functions of the ON and OFF channels of the visual system. Nature 322, 824–825.

Scott, K., Brady, R., Cravchik, A., Morozov, P., Rzhetsky, A., Zuker, C., and Axel, R. (2001). A chemosensory gene family encoding candidate gustatory and olfactory receptors in Drosophila. Cell 104, 661–673.

Simon, S.A., De Araujo, I.E., Gutierrez, R., and Nicolelis, M.A.L. (2006). The neural mechanisms of gustation: A distributed processing code. Nature Reviews Neuroscience 7, 890–901.

Sellier, M.J., Reeb, P., and Marion-Poll, F. (2011). Consumption of bitter alkaloids in Drosophila melanogaster in multiple-choice test conditions. Chemical Senses 36, 323–334.

Strother, J.A., Nern, A., and Reiser, M.B. (2014). Direct observation of on and off pathways in the drosophila visual system. Current Biology 24, 976–983.

Talay, M., Richman, E.B., Snell, N.J., Hartmann, G.G., Fisher, J.D., Sorkaç, A., Santoyo, J.F., Chou-Freed, C., Nair, N., Johnson, M., et al. (2017). Transsynaptic Mapping of Second-Order Taste Neurons in Flies by trans-Tango. Neuron 96, 783-795.e4.

Thorne, N., Chromey, C., Bray, S., and Amrein, H. (2004). Taste perception and coding in Drosophila. Current Biology 14, 1065–1079.

Vosshall, L.B., and Stocker, R.F. (2007). Molecular architecture of smell and taste in Drosophila. Annual Review of Neuroscience 30, 505–533.

Wang, Z., Singhvi, A., Kong, P., and Scott, K. (2004). Taste representations in the Drosophila brain. Cell 117, 981–991.

Weiss, L.A., Dahanukar, A., Kwon, J.Y., Banerjee, D., and Carlson, J.R. (2011). The molecular and cellular basis of bitter taste in Drosophila. Neuron 69, 258–272.

Yapici, N., Cohn, R., Schusterreiter, C., Ruta, V., and Vosshall, L.B. (2016). A Taste Circuit that Regulates Ingestion by Integrating Food and Hunger Signals. Cell 165, 715–729.

Yarmolinsky, D.A., Zuker, C.S., and Ryba, N.J.P. (2009). Common Sense about Taste: From Mammals to Insects. Cell 139, 234–244.

